# *Acidovorax temperans* polarizes T17 cells and skews neutrophil maturation to promote lung adenocarcinoma development

**DOI:** 10.1101/2022.12.20.521215

**Authors:** Joshua K. Stone, Natalia von Muhlinen, Chenran Zhang, Ana I. Robles, Eleazar Vega-Valle, Akihiko Miyanaga, Masaru Matsumoto, K. Leigh Greathouse, Tomer Cooks, Giorgio Trinchieri, Curtis C. Harris

## Abstract

Dysbiosis, or changes within the microbiome, is a common feature of solid tumors, however whether this dysbiosis directly contributes to tumor development is largely unknown. We previously characterized the lung cancer microbiome and identified *Acidovorax temperans* as enriched in tumors. In this study, we instilled *A. temperans* in an animal model driven by mutant *Kras* and *Tp53* alleles. This revealed *A. temperans* accelerates tumor development and burden through infiltration of proinflammatory cells. Neutrophils exposed to *A. temperans* displayed a mature, pro-tumorigenic genotype with increased cytokine signaling, with a global shift away from IL-1β signaling. Neutrophil to monocyte and macrophage signaling upregulated MHC II to activate CD4^+^ T cells which polarized to an IL-17A^+^ phenotype detectable in CD4^+^ and γδ populations. T17 cells shared a common gene expression program predictive of poor survival in human LUAD. These data indicate dysbiosis promotes tumor growth by modulating inflammation.

## INTRODUCTION

Lung cancer is the leading cause of cancer-specific death in the USA and worldwide (Siegel et al., 2018). Poor patient outcome is partially due to an inability to predict those patients who are likely to recur (Bray et al., 2018; Siegel et al., 2018), thus the identification and development of novel biomarkers is critical. Tobacco smoking is the predominant risk factor for lung cancer, and directly induces tumorigenesis through multiple paths, including carcinogenic metabolites, oxidative stress, and inflammation (Hecht, 2012; O’Callaghan et al., 2010).

Initial immune response to tobacco smoke is driven by IKKβ/NF-κB signaling in macrophages and an increase in proinflammatory cytokines such as IL-1β and IL-6 (Yamaguchi et al., 2012). Later stage responses see an influx of dendritic cells, neutrophils, and CD4^+^ T-cells (Bracke et al., 2006). In addition to tobacco smoking, exposure to pathogens is also believed to play a pro-inflammatory role by creating a local environment primed for oncogenesis. Infection with *Mycobacterium tuberculosis* has been linked to an increased risk for lung cancer development (Brenner et al., 2001; Engels et al., 2009; Shiels et al., 2011), possibly through increased infiltration of proinflammatory cells such as neutrophils (Kroon et al., 2018).

Recently, the native microbiota was identified as a key regulator of immune function in an autochthonous mouse model of lung adenocarcinoma (LUAD). Mice kept in specific pathogen-free (SPF) conditions developed more and larger tumors compared to those kept in germ-free (GF) conditions. The presence of bacteria resulted in a proinflammatory microenvironment, characterized by recruitment of IL-1β-secreting alveolar macrophages, which in turn activated IL17-secreting γδ T-cells, finally recruiting large numbers of neutrophils to the tumor, indicating a role for bacteria in tumor growth (Jin et al., 2019a). In lung cancer patients, lower airway microbes were associated with infiltration of T_H_17 cells and neutrophils (Segal et al., 2016). These results suggest an important proinflammatory role for the microbiome in the development of lung cancer.

Our group recently showed that the lung microbiome undergoes dysbiosis in cancer patients (Greathouse et al., 2018). We identified the Gram-negative *Acidovorax* genus as differentially abundant between normal and tumor tissue as well as between lung adenocarcinoma and squamous cell carcinoma. Furthermore, *Acidovorax* abundance was linked to smoking status and *TP53* mutations (Greathouse et al., 2018), supported by the detection of *Acidovorax* in tobacco cigarettes (Malayil et al., 2020). Full-length 16S rRNA gene sequencing as well as fluorescence *in situ* hybridization identified *Acidovorax temperans* in patient tumors. Our findings were later confirmed by multiple studies, which detected *Acidovorax* 16S signal in both tumor tissue and patient sputum (Jin et al., 2019b; Leng et al., 2021; Shimizu et al., 2022; Yu et al., 2016).

A central question that has emerged from these sequencing-based studies of the microbiome is whether dysbiosis plays a causative or correlative role in tumorigenesis, i.e., is the microbiome a driver or passenger microenvironmental factor? The Jacks lab (Jin et al., 2019a) began to address this question by linking commensal bacteria, inflammation, and LUAD growth. However, the role dysbiosis may play in promoting tumorigenesis is largely unknown.

To answer this question, we repeatedly instilled *A. temperans* as a model for dysbiosis into an autochthonous mouse model of LUAD driven by K-ras and Tp53 mutations to mimic the effects of chronic smoking. We found a driver role for *A. temperans* in tumor growth whereby bacterial instillation increased tumor growth through inflammation, primarily driven by neutrophils, macrophages, and CD4^+^ T cells. This proinflammatory response suggests dysbiosis in the presence of driver mutations in epithelial cells is sufficient to promote a tumorigenic microenvironment.

## RESULTS

### *Acidovorax temperans* exposure accelerates tumor development in an autochthonous LUAD mouse model

Previous research developed a mutant K-ras and Tp53-driven LUAD mouse model (KP) under the Cre-lox system, resulting in endogenous tumor development reflective of human LUAD (Jackson et al., 2005; Jackson et al., 2001). Our group identified *Acidovorax temperans* as enriched in lung cancer which led us to hypothesize that *A. temperans* may play a functional role in lung cancer development (Greathouse et al., 2018). To determine if repeated bacterial exposure, as experienced in chronic tobacco smoking, would result in increased tumor growth, we administrated six biweekly intranasal instillations of PBS (sham) or *A. temperans* in KP mice following Ad-cre instillation (Figure 1A).

**Fig. 1.**
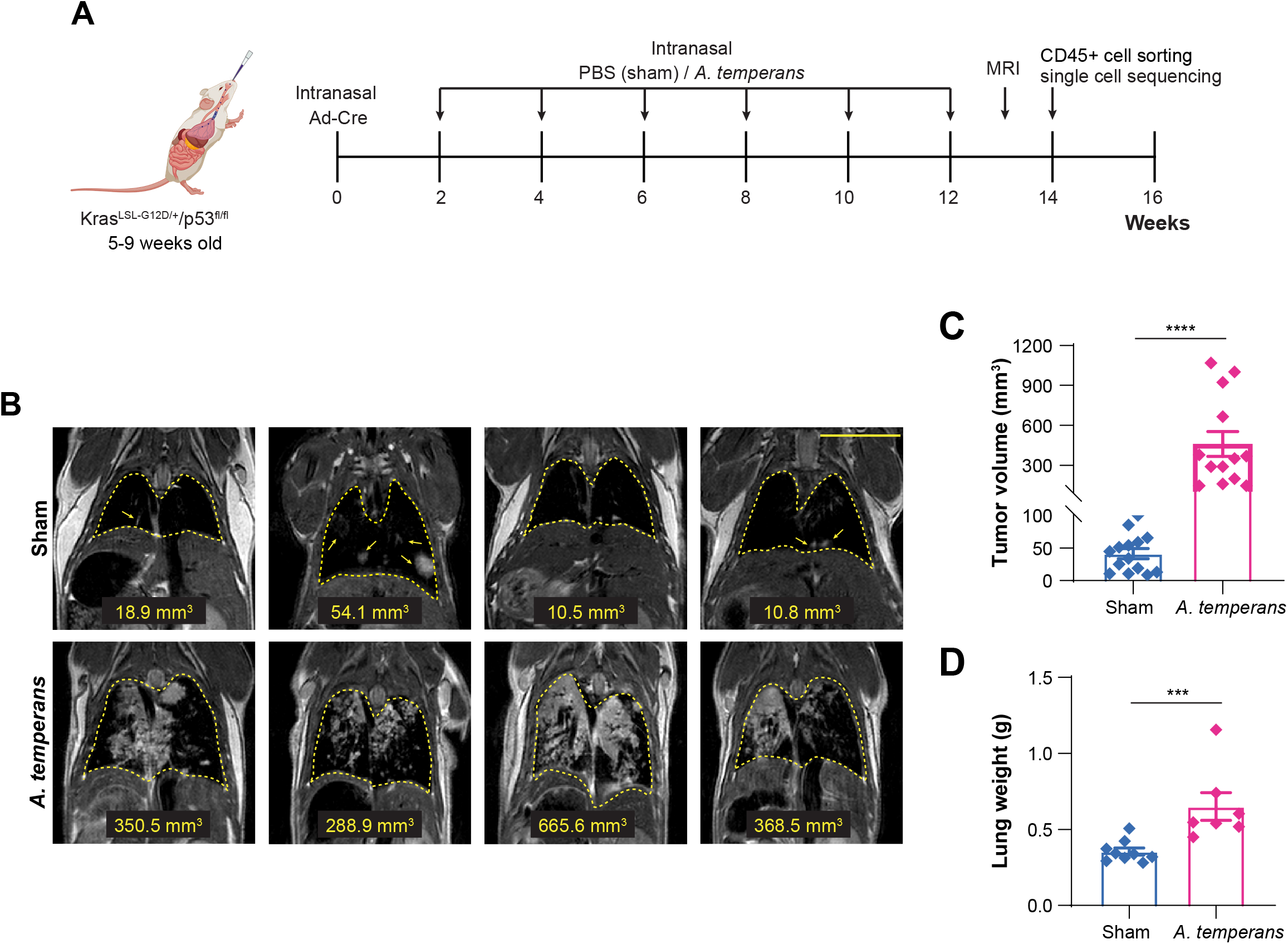
*Acidovorax temperans* accelerates tumor growth in a mouse model of lung adenocarcinoma. (A) Experimental timeline including bacterial dosage schedule. (B) MRI images of sham (1X PBS) (top) and *A. temperans* (bottom) instilled mice. (C) MRI quantification of tumor volume in sham (n = 14) and *A. temperans* (n = 13) instilled mice. (D) Quantification of lung weight in sham (n = 8) and *A. temperans* (n = 7) instilled mice. Data presented as mean ± SEM, *** *p* < 0.001, **** *p* < 0.0001.

Using non-invasive magnetic resonance imaging (MRI), we measured tumor development 13 weeks post Ad-cre instillation and then sacrificed mice one week later. We found that *A. temperans* instilled mice had visibly larger nodules compared to those instilled with sham by MRI and quantification demonstrated an increase in tumor volume (Figures 1B, 1C). Consistent with these results, total lung weight was also increased in *A. temperans* mice (Figure 1D). Taken together, these results revealed that repeated exposure to *A. temperans* could accelerate lung tumor development in the presence of oncogenic K-ras and Tp53 mutations.

### Immune cell infiltration within the tumor microenvironment is altered by *A. temperans*

Previous studies have demonstrated an important role for a proinflammatory tumor microenvironment in KP LUAD development, with dendritic cells (Maier et al., 2020), macrophages (Cortez-Retamozo et al., 2012), neutrophils (Ancey et al., 2021; Engblom et al., 2017; Faget et al., 2017; Simoncello et al., 2022), and T cells (Horton et al., 2021; Jin et al., 2019a) all implicated in the etiology of this animal model. Considering the overlap of these cell types with those involved in bacterial response, we hypothesized that dysbiosis may accelerate tumor growth by altering the immune microenvironment.

To test this hypothesis, we dissociated lung tissue from four sham and four *A. temperans* instilled KP mice and isolated the CD45^+^ fraction by FACS (Figure 1A). We then performed droplet-capture single cell RNA sequencing (scRNA-seq), which returned 25,477 total CD45^+^ cells after filtering. These cells divided into 11 major cell types: monocytes, macrophages, monocyte-derived dendritic cells (MoDC), alveolar macrophages (AMs), conventional dendritic cells (cDC), plasmacytoid dendritic cells (pDC), neutrophils, B cells, plasma cells, NK cells, and T cells (Figure 2A). Cell types largely overlapped between sham and *A. temperans* mice, although cell proportions showed largest relative shifts from monocyte-high in sham to increased macrophages, AMs, neutrophils, and T cells in *A. temperans* mice (Figures 2A, 2B). We identified these clusters through three methods, by comparison to the ImmGen database (Figure 2C) (Heng et al., 2008), canonical gene markers (Figure 2D), and top differentially expressed genes (Figure 2E, Supplementary Table S1). Collectively, this demonstrates *A. temperans* alters the immune compartment of the tumor microenvironment in KP mice.

**Fig. 2.**
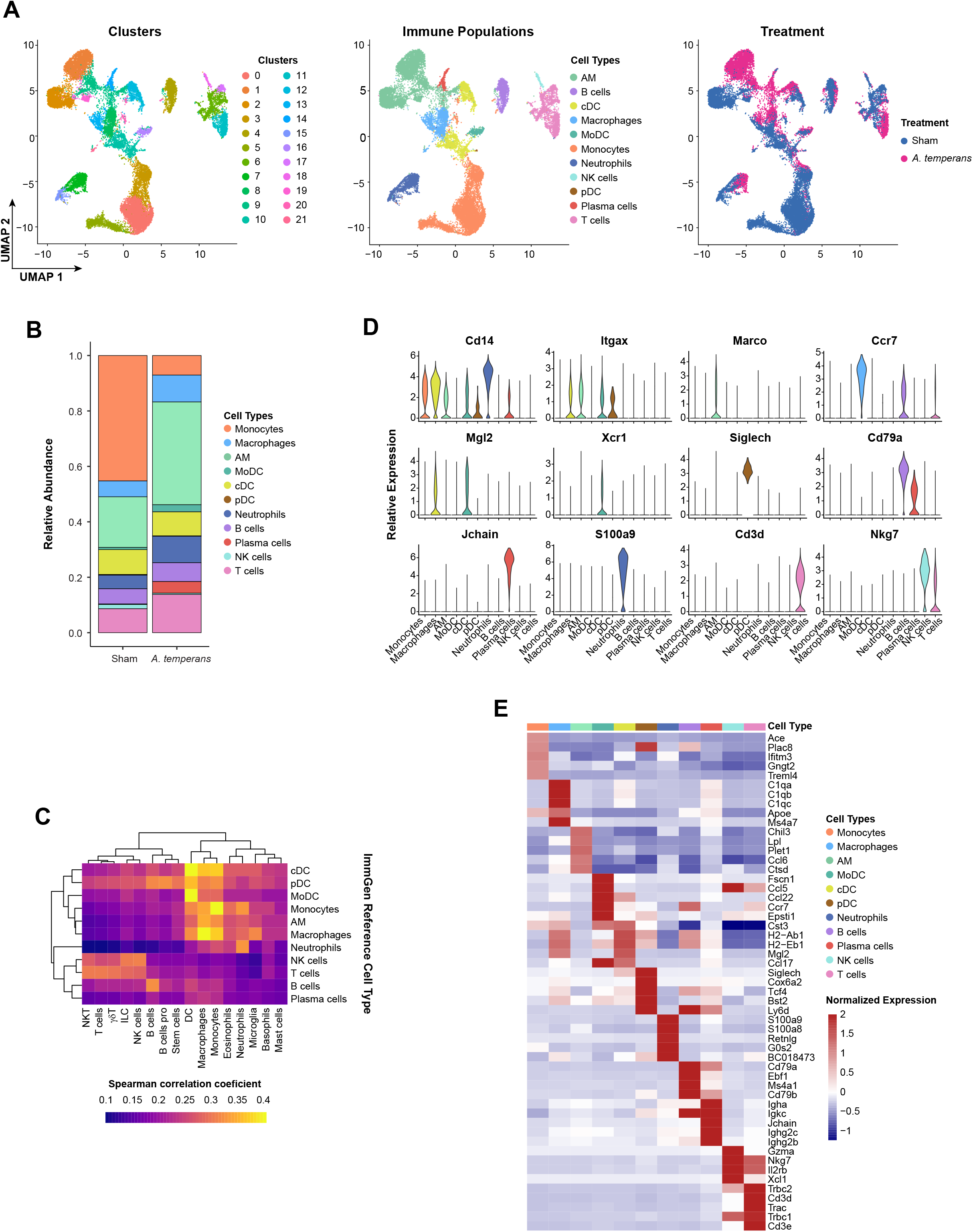
*A. temperans* alters the immune microenvironment in lung adenocarcinoma. (A) UMAP plots of scRNA-seq cell clusters (left), immune cell types (center), and treatment groups (right). (B) Barplot of the relative abundance for each cell type by treatment group. (C) Heatmap of ImmGen-based cell type identification. Color scale indicates positive Spearman’s correlation coefficient. (D) Violin plots for marker genes associated with the different cell types. (E) Heatmap of the top five differentially expressed genes for each cell type.

### Lung macrophages upregulate MHC class II in response to *A. temperans*

Myeloid cells are the first cells to respond to bacterial lung infections and often secrete proinflammatory cytokines linked to tumor development, leading us to first characterize this compartment. We identified 13 subclusters corresponding to monocyte, macrophages, and dendritic cells (MoMaDCs) (Figures 3A, 3B). Overall, we identified two clusters of naïve monocytes (*Cd14+; Fcgr3-*, CD16), three activated monocyte clusters (Act Mono; *Cd14+, Fcgr3+*), one cycling monocyte cluster (*Mki67, Stmn1, Top2a*), three macrophage clusters (*Cd68*), and four DC clusters (*Syngr2*) (Figure 3C, Supplemental Table S2). Within the macrophages, we identified two clusters of tumor-associated macrophages (TAMs; *Fcgr2b, Ccl4, Trem2*) (Zhang et al., 2020) and one enriched in complement genes (*C4b, Cfp, C1qb*). Within the DCs, we identified MoDCs (*Ccl5, Ccr7, Fscn1*), conventional DCs clusters cDC1 (*Clec9a, Itgae, Xcr1*) (Minoda et al., 2017) and cDC2 (*Mgl2, Irf4*), and plasmacytoid DCs (*Bst2, Pacsin1, Siglech*) (Figure 3C, Supplementary Table S2) (Abbas et al., 2020). Cell type identification was confirmed by comparison to the ImmGen database (Figure 3D). We then asked if developmental trajectory followed the conventional route from circulating monocytes to macrophages and DCs. First removing the cDC and pDC clusters as these cell types are not monocyte derived, this analysis revealed that the TAMs and MoDCs were the latest in pseudotime, representing the most terminally differentiated cell states (Figures 3E, 3F).

**Fig. 3.**
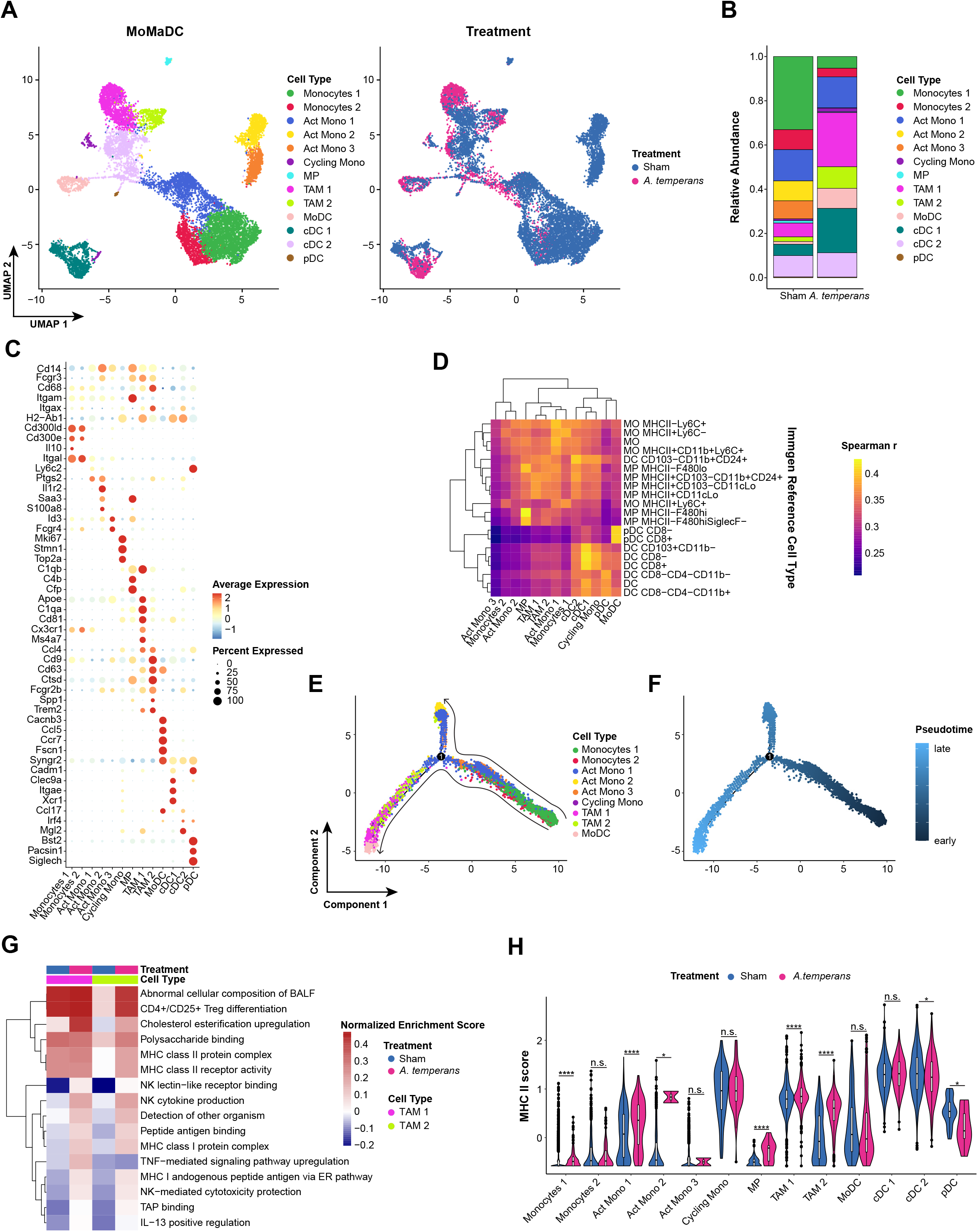
TAMs are expanded and upregulate MHC II in response to *A. temperans*. (A) UMAP plots of monocytes, macrophages, and dendritic cells (MoMaDCs) cell types (left) and treatment groups (right). (B) Barplot of the relative abundance for each cell type by treatment group. (C) Dotplot of marker genes for each cell type. (D) Heatmap of ImmGen-based cell type identification. Color scale indicates positive Spearman’s correlation coefficient. (E) UMAP plots of trajectory analysis of monocyte-derived cells. (F) UMAP plots of pseudotime projection of monocyte-derived cells. (G) Single-sample GSEA (ssGSEA) heatmap of average normalized enrichment scores for both TAM clusters divided by treatment group. (H) Comparison of average expression of each MHC II component gene (*H2-Aa, -Ab1, -DMa, - DMb1, -DMb2, -Eb1, -Eb2, -Oa, -Ob*) by treatment for each cell type. Data presented as median value plus quartiles for boxplots, n.s. not significant, * *p* < 0.05, **** *p* < 0.0001.

Macrophages have traditionally been classified as M1 or M2, with classically activated M1 macrophages inducing inflammation against pathogens and tumor cells while M2 macrophages are immunosuppressive. Neither TAM cluster showed expression of M1 or M2 macrophage markers (Figure S1A). To better understand differences in their function, we examined changes in TAM gene expression with single-sample gene set enrichment analysis (ssGSEA) (Borcherding et al., 2021). These clusters were differentiated by cholesterol esterification upregulation in TAM-1 in both treatment groups while TNF signaling was specifically upregulated in TAM-1 in response to *A. temperans* (Figure 3G). Both TAM-1 and TAM-2 clusters were highly enriched for both MHC class I and II antigen presentation in response to *A. temperans* instillation. To determine if MHC upregulation was consistent among monocytes and DCs, we calculated signature scores for all MHC class I and II genes. This revealed *A. temperans* caused upregulation of MHC I in macrophages and DCs, but only macrophages displayed consistent upregulation of MHC II, with TAM-2 cells having the greatest relative increase (Figures 3H, S1B, S1C). Similarly, analysis of the alveolar macrophage (AM) compartment revealed low expression of M1/M2 genes while MHC I and II were both broadly upregulated across AMs in response to *A. temperans* (Figure S2, Supplementary Table S3). Together, these results indicate that dysbiosis induces a broad MHC II response across lung macrophages, potentially contributing to increased tumor growth through CD4^+^ T cell activation.

### *A. temperans* increases tumor-associated neutrophils which express antimicrobial gene programs

Neutrophils are the most abundant immune cell type in human NSCLC and KP mice (Faget et al., 2017; Kargl et al., 2017) and have dual function in cancer development and infection response, particularly secretion of proinflammatory cytokines, ROS production, and immunosuppression (Hedrick and Malanchi, 2022). These factors implicate neutrophils as essential mediators of early tumor development, therefore we asked how *A. temperans* altered expression and function of neutrophils in KP mice.

We identified four clusters of neutrophils, with sham mice having greater proportions of clusters C1 and C2 while *A. temperans* instilled mice greatly increased the cell numbers in clusters C0 and C3 (Figures 4A, 4B). Markers associated with circulating neutrophils (*Sell^hl^*, CD62L; *Cxcr4^lo^*) were higher in clusters C1 and C2 while markers linked to increased effector function (*Icam1*), immunosuppression (*Cd274*, PD-L1), and tumor-promotion (*Siglecf*) were higher in clusters C0 and C3; these expression patterns generally corresponded with treatment group (Figure 4C, Supplementary Table S4) (Pfirschke et al., 2020; Woodfin et al., 2016). To verify this change in marker gene expression from control to bacterial exposure, we performed trajectory analysis on the scRNA-seq clusters. This revealed C1 was earliest and C3 latest in pseudotime (Figure 4D), consistent with expression of these marker genes.

**Fig. 4.**
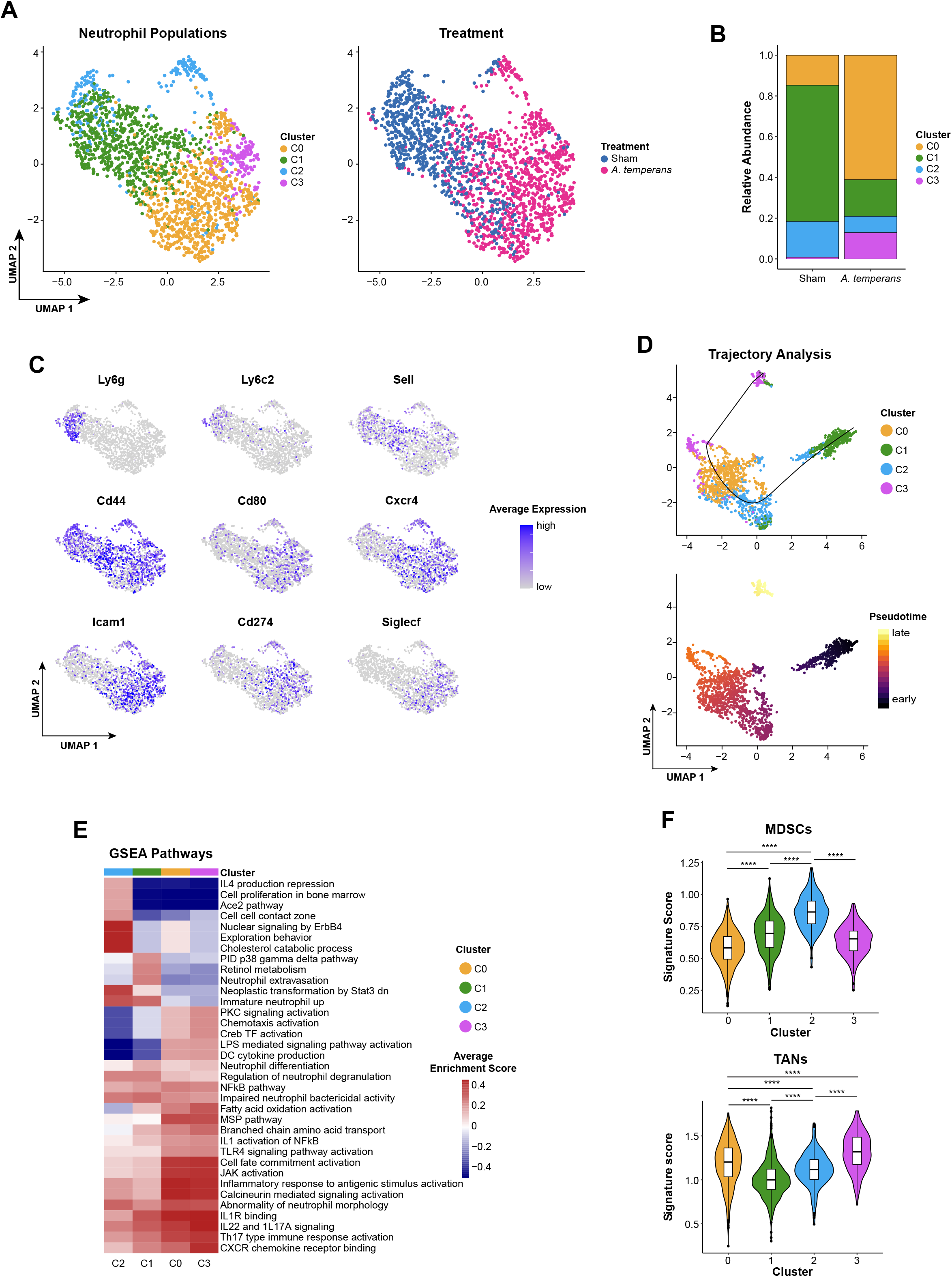
Tumor-associated neutrophils are associated with increased anti-bacterial function. (A) UMAP plots of neutrophil clusters (left) and treatment groups (right). (B) Barplot of the relative abundance for each cluster by treatment group. (C) Density plots of marker gene expression. (D) UMAP plots of trajectory analysis (top) and pseudotime (bottom) of neutrophil clusters. (E) ssGSEA heatmap of average normalized enrichment scores for each neutrophil cluster. (F) Gene signature scores for granulocytic myeloid-derived suppressor cells (MDSCs) (top) and tumor associated neutrophils (TANs). Accession numbers for gene signatures: GSE139125 (MDSCs) and GSE118245 (TANs) (Alshetaiwi et al., 2020; Mollaoglu et al., 2018). Boxplots indicate median and quartile scores. **** *p* < 0.0001

To further understand transcriptional differences between sham and *A. temperans-* associated neutrophils, we performed ssGSEA, which suggested C2 represented the most immature cell state, with high enrichment scores for cell proliferation in bone marrow, immature neutrophils, and cholesterol catabolism, important for neutrophil development and release from the bone marrow (Figure 4E) (Cavalcanti et al., 2007). C1 pathways were enriched for neutrophil extravasation while activated, effector functions such migration, chemotaxis, and bacterial response and killing, were primarily associated with C0 and C3, underlined by an LPS response which activated cytokine production.

We then asked if these changes reflected gene expression profiles of immunosuppressive granulocytes in cancer. Granulocytic myeloid-derived suppressor cells (MDSCs) are key suppressors of T cells and phenotypically analogous to neutrophils, but genetically distinct. We compared a gene signature of MDSCs to our neutrophil clusters (Alshetaiwi et al., 2020), which revealed C2 had the highest expression of the MDSC signature (Figure 4F). Tumor-associated neutrophils (TANs) are also an immunosuppressive population required for tumor progression and metastasis (Giese et al., 2019), and comparison of a TAN gene signature revealed C3 had the highest expression of this signature (Figure 4F) (Mollaoglu et al., 2018). Collectively, these results demonstrate that dysbiosis response is a key programming event for tumor-associated neutrophils, suggesting that these neutrophils, while being responsible for clearing bacteria from the lungs, alter the tumor microenvironment.

### *A. temperans* robustly induces T_H_17 polarization

Having demonstrated large proinflammatory changes in the myeloid compartment driven by MHC upregulation in MoMaDCs, we then examined if these changes were reflected in T cells, as well. We identified a total of 12 T cell types and sham mice had higher proportions of naïve T cells while *A. temperans* mice showed greater CD4^+^ effector populations (Figures 5A, 5B, Supplemental Table S5). These effector populations included follicular helper T cells (Tfh; *Cd200, Izumo1r, Slamf6*) (Künzli et al., 2020), Tregs (*Foxp3, Ikzf2, Ctla4*), Th17 (*Il17a, Tmem176a, Tmem176b*) (Ciofani et al., 2012), and Th1 (*Ccr2, Ifng*). We found three clusters of cells which did not express either *Cd4* or *Cd8a* (Figure 5C), which corresponded to γδ T cells (*Tcrg-C1, Trdc*), double negative (DN) naïve (*Ccr7, Lef1, Sell, Tcf7*), and a DN Treg-like population (*Areg, Gata3, Il1rl1*) (Miragaia et al., 2019). Marker genes for various known DN T subtypes were also not expressed in DN naïve and Treg-like cells, preventing further identification of these cells (Figure S3).

**Fig. 5.**
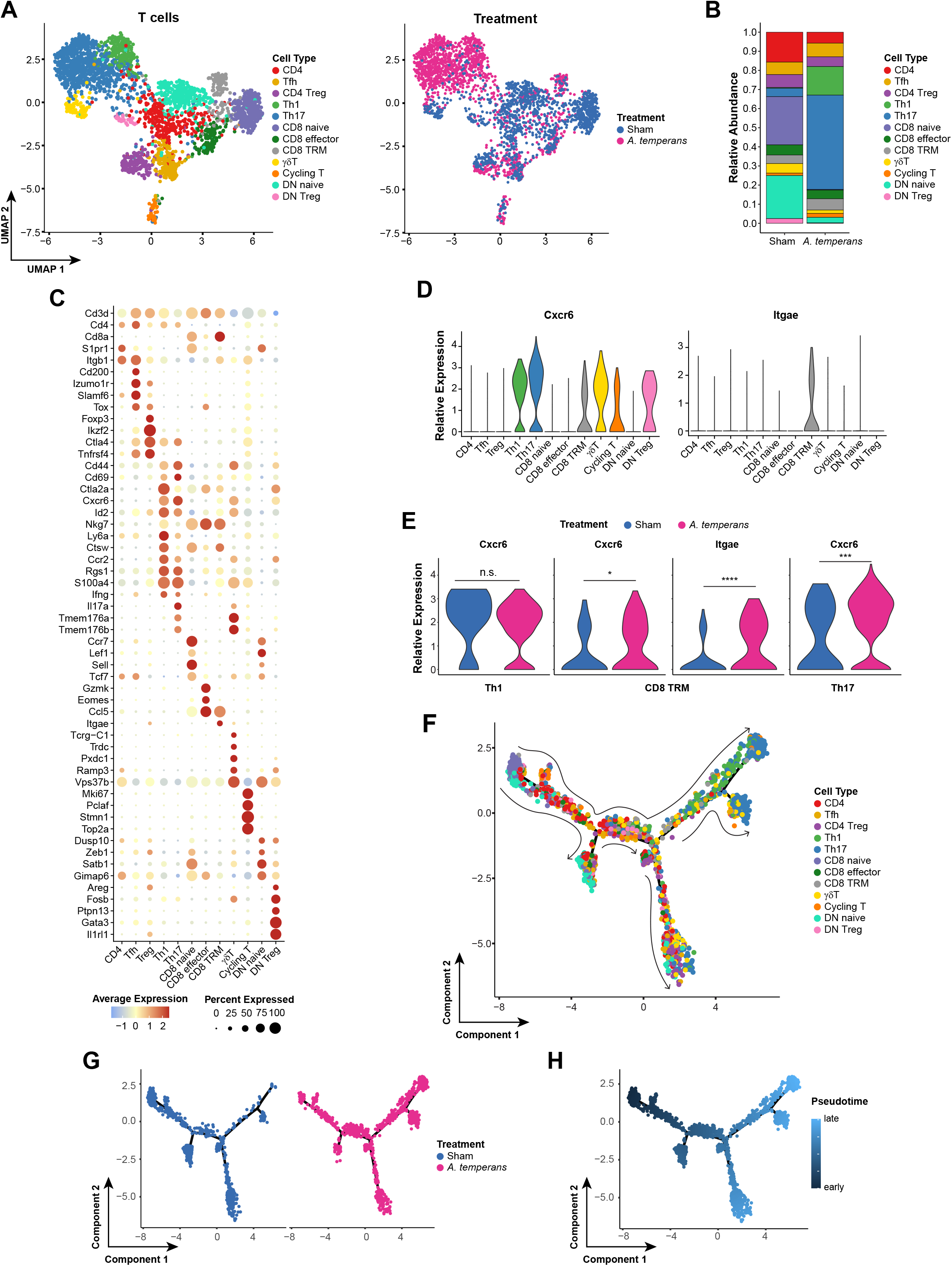
T cells are terminally polarized to a Th17 genotype. (A) UMAP plots of T cell subtypes (left) and treatment groups (right). (B) Barplot of the relative abundance for each cell type by treatment group. (C) Dotplot of marker genes for each cell type. (D) Density plot for marker genes. (E) Expression level of tissue residency marker genes by cell type. (F) Expression level of tissue residency markers for Th1 (left), CD8 TRM (center), and Th17 (right) by treatment group. (G and H) UMAP plots of trajectory analysis by (G) cell type (left) and pseudotime projection (right) and (H) by treatment group. n.s. not significant, * *p* < 0.05, *** *p* < 0.001, **** *p* < 0.0001.

T cells from *A. temperans* mice also showed greater expression of tissue residency markers *Cxcr6* and *Itgae* (CD103) in CD4^+^ and CD8^+^ T cells (Figures 5D, 5E). As tissue residency is associated with effector function, we then asked if these effector CD4^+^ cells represented a terminal cell state. Trajectory analysis which revealed naïve CD8 and DN cells were earliest in pseudotime, while Th1 and Th17 cells were latest (Figures 5F – 5H). The similarity of marker genes between the Th1 and Th17 cells, combined with high expression of *Cxcr6* in Th1 and Th17 cells, suggested that effector CD4^+^ T cells acquire a tissue residency phenotype prior to polarization. In support of this hypothesis, *A. temperans* T cells were consistently later in pseudotime than sham (Figure 5G), suggesting that dysbiosis is a key factor for establishing CD4^+^ T cell lung residency and subsequent Th1/Th17 polarization.

### *A. temperans* induces specific IL-17 and broad IFN-γ response in T cells

Previous data examining murine colonic effector T cells suggested that T cell phenotype was shaped by response to specific pathogens (Kiner et al., 2021). We asked if the T cell polarization induced by *A. temperans* was specific to this species or was consistent with a more general microbial dysbiosis. We compared gene signatures of murine infections with either *Citrobacter rodentium* (T_H_17 response) or *Salmonella enterica* serovar Typhimurium (IFN-γ response), both Gram-negative species (Kiner et al., 2021). Expression of the *C. rodentium* signature was predominantly found in our Th1, Th17, and γδ T clusters (Figures S4A, S4B). Although the *Salmonella* Typhimurium signature was also highest in Th1 and Th17 clusters, we observed consistently high expression throughout our dataset, but upregulated in *A. temperans* mice compared to sham overall (Figures S4C, S4D). These data suggest that the Th17 cell cluster we observe is not specific to *A. temperans;* however, the combination of general IFN-γ and specific T_H_17 polarization may represent a unique inflammatory response to this species.

Based on the widespread upregulation of the *Salmonella* Typhimurium gene signature, we asked if both *Il17a* and *Ifng* were upregulated in response to *A. temperans*. Our results showed *Il17a* and its transcription factor *Rorc* were largely restricted to Th17 and γδ T cells while *Ifng* and its transcription factor *Stat4* were highly expressed in non-naïve T cells (Figures S5A, S5B). Overall, most T cell subtypes expressed *Ifng*, with nearly half of Th1 and a third of Th17 cells positive for this transcript and expression was elevated in response to *A. temperans* (Figures S5C – S5E). Within Th17 cells, a subpopulation was double positive for *Il17a* and *Ifng* (Figures S5F, S5G), a highly inflammatory cell state increased in smokers (Xu et al., 2019). These data suggest that *A. temperans* alters the immune microenvironment through multiple signaling pathways which culminate in *Il17a+/Ifng+* T cells to greatly increase inflammation.

### A conserved gene signature in T17 cells induced by dysbiosis is predictive of poor survival in LUAD

In addition to Th17 cells, approximately 40% of γδ T cells also expressed *Il17a* (Figure 5C). Examining every T cell cluster revealed that most *Il17a+* cells, regardless of cell type, were from *A. temperans* mice (Figure 6A). Given the high percentage of *Il17a+* cells in Th17 and γδ T cells, we hypothesized that gene expression may be similar in both clusters which could then identify a gene set important for IL-17 polarization in pan T cell subtypes (T17). To test this, we combined these two clusters and calculated differentially expressed genes against all other T cells. The resulting T17 signature was upregulated in *Il17a+* cells in both Th17 and γδ T cells (Figures 6B, 6C, Supplemental Table S6). Examination of the T17 signature by cell type showed highest expression in Th17 and γδ T cells, as expected, but also DN Treg, cycling T, and Th1 cells (Figure 6D). Interestingly, this signature score increased during T cell differentiation in multiple subtypes, including CD4^+^, CD8^+^, and DN T cells (Figure 6D). As average expression of the T17 signature was higher in *A. temperans* mice regardless of T cell subtype (Figure 6E), these data suggested repeated exposure to *A. temperans* induces a T17 polarization in multiple T cell subtypes.

**Fig. 6.**
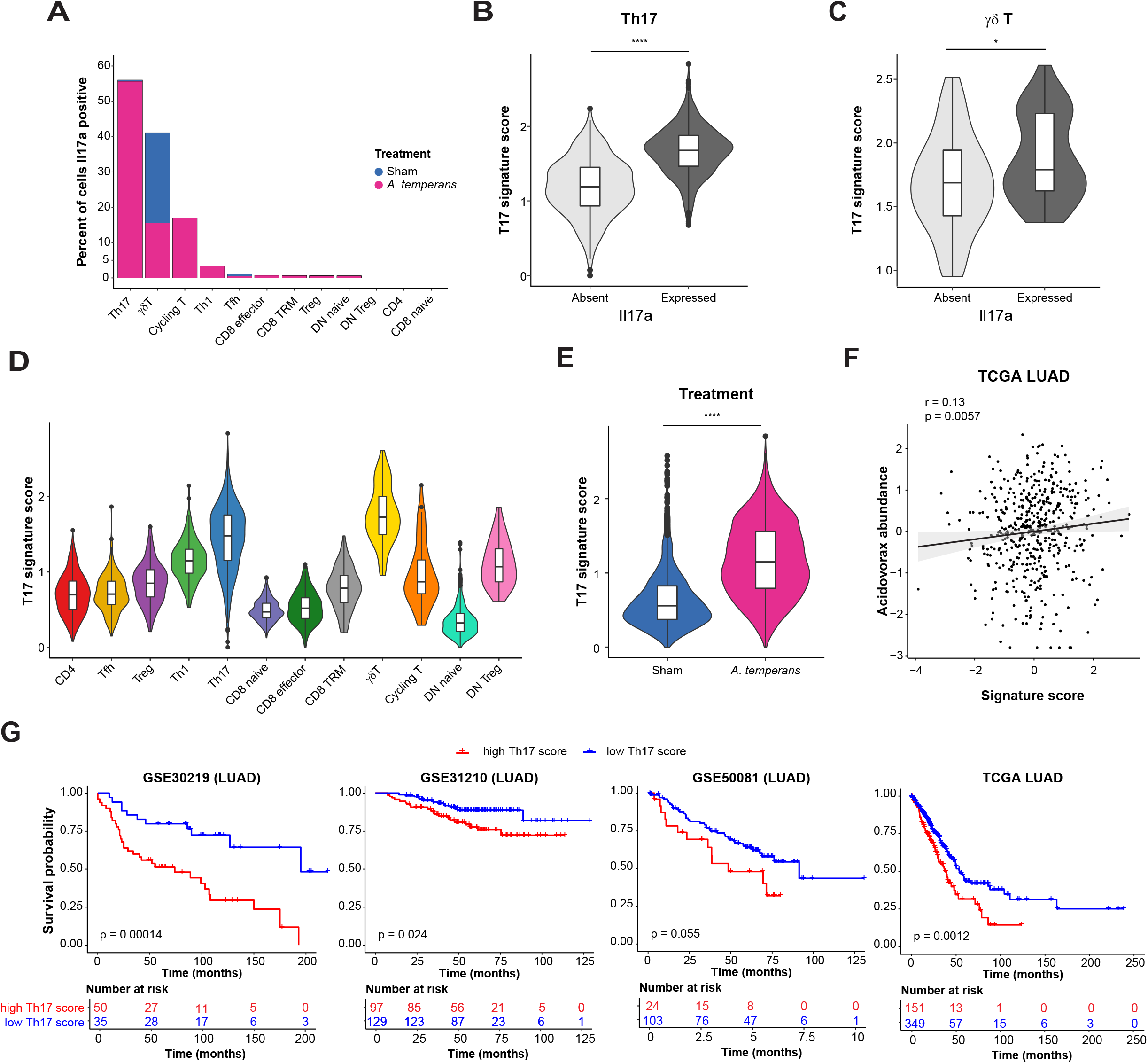
A pan T17 gene signature is predictive of poor prognosis in human LUAD. (A) Percent of cells *Il17a* positive per T cell subtype and treatment group. (B and C) Expression of a pan T17 cell gene signature in *Il17a* negative (absent) and positive (expressed) cells within the Th17 (B) and γδ T (C) clusters. (D and E) Expression of a pan T17 cell gene signature in all cells within each T cell subtype (D) and by treatment group (E). (F) Correlation of *Acidovorax* abundance (genus level) and the murine pan T17 gene signature expression within the TCGA LUAD dataset (Greathouse et al., 2018). (G) Kaplan-Meier curves for survival property within four human LUAD cohorts – GSE30219, GSE31210, GSE50081, and TCGA (Collisson et al., 2014; Der et al., 2014; Okayama et al., 2012; Rousseaux et al., 2013).

We then asked if this T17 gene signature was important in human lung cancer. Leveraging the metatranscriptomics data that we had previously generated using TCGA LUAD (Greathouse et al., 2018), we examined the association of the T17 signature with *Acidovorax* exposure in these patients. The T17 signature score was positively correlated with *Acidovorax* abundance (Figure 6F), suggesting microbial dysbiosis also results in T17 polarization in human LUAD. Next, we asked if high expression of the T17 signature was predictive of patient survival. We stratified patients by low or high expression of the T17 signature score in four cohorts of LUAD patients, including TCGA. This stratification revealed high expression of the T17 gene signature was a poor prognostic in LUAD (Figure 6G). These results suggest T17 polarization, regardless of T cell receptor type, accelerates tumor development and results in worse survival in patients.

### Cell-cell signaling switches from IL-1 β driven to broad proinflammatory activation in response to *A. temperans*

We then investigated cell-cell communication to determine the potential mechanism of *A. temperans*-mediated LUAD progression. Examination of cytokines within all cell types revealed immune cell-of-origin for multiple cytokines previously implicated in KP mouse etiology: IL-1β (neutrophils), IL-23 (neutrophils), and IL-17 (T cells) (Figure 7A). In contrast to previous results, IL-22 was not detected, and AMs were not a major source of IL-1β or IL-23 (Jin et al., 2019a), suggesting that introduction of an external bacterial species dramatically alters immune cell signaling in KP mice.

**Fig. 7.**
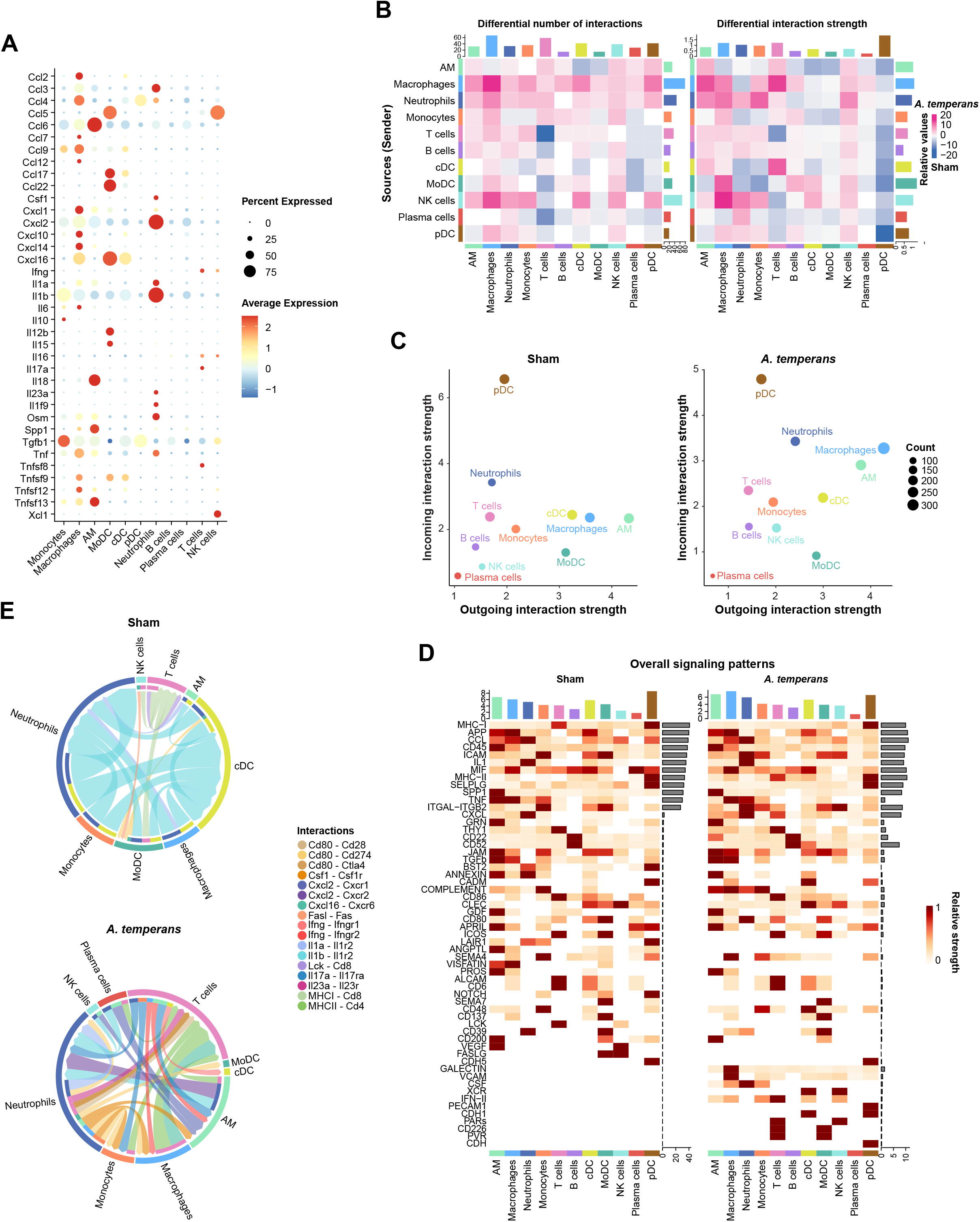
Cell-cell communication demonstrates *A. temperans* induces robust signaling between neutrophils, macrophages, and T cells. (A) Dotplot of cytokine expression by cell type. (B) Heatmap depicting differential interactions as calculated by expression of ligand in outgoing cell type and its cognate receptor in the incoming cell type, with total number (left) and interaction strength (right). Interactions enriched in sham mice are indicated by blue and those enriched in *A. temperans* mice indicated by pink. (C) Cross-referenced incoming and outgoing interaction strength for each cell type in sham (left) and *A. temperans* (right) mice. (D) Heatmap of overall signaling patterns by cell signaling pathway for each cell type in sham (left) and *A. temperans* (right) mice. (F) Chord diagrams showing pathways in significantly enriched in Sham (top) and *A. temperans* mice (bottom). Chord diameter indicates aggregate expression of the ligand and receptor, arrow indicates direction from sender to receiver population, outer rings indicate sender cell type, inner rings indicate receiver cell type, and links are colored by interaction pairing. Individual chords are colored by secreting cell type and arrows indicate receptor cell type.

We then compared cell-cell communication by examining combined changes in expression of ligand-receptor pairs in sham and *A. temperans* mice. Examination of the total signaling strength (combined incoming and outgoing signal) revealed a specific increase in number of interactions among neutrophils, AMs, macrophages, and T cells in response to *A. temperans* (Figures 7B, 7C, Supplementary Tables S7, S8). Sham mice were enriched for BST2, LAIR1, FASLG, IL1B, and VEGF ligand-receptor signaling while *A. temperans* mice were enriched for a MHC class I to class II switch, CXCL, IFN type II, and CSF signaling (Figure S6). In-depth analysis of key ligand-receptor pathways indicated an increase in *Csf1-Csf1r* signaling between neutrophils and monocytes/macrophages; *Ifng/Ifngr1/2* between T cells and monocytes, macrophages, and AMs; *Cxcl2-Cxcr1/2* between neutrophils and AMs; and a strong increase in *MHC II-CD4* between macrophages, cDC, and T cells (Figures 7D, 7E). *Il17a-Il17ra* interactions were not predicted due to their expression levels below threshold in sham mice which prevented fold differences from being calculated, demonstrating specific expression in non-γδ T cells in *A. temperans* mice.

Overall, these results indicate that dysbiosis causes massive inflammation in the KP mouse model of lung cancer, with mature, tumor-associated neutrophils secrete Csf1 to promote differentiation of monocytes to macrophages, which then strongly upregulate MHC class II to stimulate T cells into T17 polarization. This proinflammatory cell infiltration provides mechanistic insights into how dysbiosis alters and promotes lung cancer development, which may have implications for smoking-related tumorigenesis.

## DISCUSSION

Previous studies have demonstrated lung and other cancers feature dysbiotic microbiomes, but a central question is if this dysbiosis contributes to tumor growth (Ramírez-Labrada et al., 2020; Xavier et al., 2020). We hypothesized that changes to the microbiome in lung cancer patients likely resulted from repeated exposures, through smoking and/or oral microaspirations (Dickson et al., 2017). To mimic these multiple exposures, we repeatedly instilled *A. temperans*, which we previously identified as associated with smoking and *TP53* mutations in human lung cancer (Greathouse et al., 2018), in the KP mouse model of LUAD. This revealed that *A. temperans* increased tumor burden and development, partially through large proinflammatory changes to the tumor microenvironment, driven by tumor-associated neutrophils, MHC class II expressing macrophages, and T17 cells.

Neutrophils and AMs likely form the first-line response to *A. temperans* instillation and secretion of IL-1β and IL-23 from these cells were previously found to be critical for activation of γδ T17 cells in KP mice, which then recruited neutrophils to the tumor site (Jin et al., 2019a). We also found *Il1b* was expressed by monocytes and macrophages but observed that the primary source of *Il1b* was neutrophils themselves, which was secreted in an autocrine loop or to cDCs in sham mice. These differences are likely directly attributable to the instillation of *A. temperans* to the lung, as Jin et al. (Jin et al., 2019a) compared KP mice kept in SPF and GF conditions, identifying γδ T17 cells as the key subpopulation driving inflammation. In keeping both sham and *A. temperans* instilled mice in SPF conditions, our data suggests that rather it is neutrophils that are the primary drivers of inflammation following dysbiosis. Using an orthotopic mouse model with cancer cells derived from KP mice, Tsay et al. (Tsay et al., 2021) also found a similar pathway in response to the oral commensal *Veillonella parvula*. Following *V. parvula* instillation in the lungs, they reported an increase in T_H_17 and γδ T17 cells which attracted large infiltration of neutrophils. Collectively, these results indicate a key role for bacterially induced T17 cells and neutrophils in the development of murine LUAD.

Mature, SiglecF^high^ neutrophils are characterized by a long intratumoral half-life (3-5 days), lack of proliferation, increased ROS production, and promotion of tumor cell proliferation (Engblom et al., 2017; Pfirschke et al., 2020). The source of these cells is unknown and SiglecF^high^ neutrophils are not found in the bone marrow, circulation, or spleen (Pfirschke et al., 2020). Instead, it is thought that these cells complete maturation in the lungs after infiltrating the tumor site. Similarly, immature neutrophils are found in the circulation of inflammation patients and undergo maturation at the site of *Staphylococcus aureus* infections (Coffelt et al., 2016; Kim et al., 2011). *Siglecf* expression was increased in response to *A. temperans* instillation and associated with strong immune function, including LPS response and IL-17 signaling. Our data suggest repeated exposure to even transient dysbiosis recruits immature neutrophils to the lung where they undergo localized maturation and persistence for tumor promotion, although further studies are needed to test this hypothesis.

Our data have several aspects that are directly relatable to human lung cancer. First, we previously identified *Acidovorax* spp. enriched in smokers and hypothesized that tobacco smoke was a possible means for introducing *Acidovorax* into the lungs (Greathouse et al., 2018). Smoking gradually reduces epithelial barrier function, potentially allowing bacteria such as *Acidovorax* direct access to tumors as a route for inducing dysbiosis in multiple pulmonary diseases (Huang and Shi, 2019). *Acidovorax* spp. have been detected in cigarettes and possess genes for catabolism of smoking-associated hydrocarbons (Malayil et al., 2020; Singleton et al., 2009). Smoking also results in a similar inflammatory pathway to what we identified in *A. temperans* mice, potentially a response to the large amounts of LPS contained in cigarettes (Bracke et al., 2006; Hasday et al., 1999; Larsson et al., 2008; Yamaguchi et al., 2012). LPS by itself is capable of accelerating lung cancer growth in vivo, partially through macrophage infiltration and activation of NF-κB and STAT3 signaling (Melkamu et al., 2013), providing an additional route for smoking-induced oncogenesis. Second, neutrophils are the most abundant immune cell found in human lung cancer and high infiltration is generally associated with both poor prognosis and resistance to various therapies (Coffelt et al., 2016; Kargl et al., 2017; Rapoport et al., 2020). Tumor-associated neutrophils, characterized by an activated phenotype of CD62L^low^ (*SELL*)/CD54^high^ (*ICAM1*) (comparable to our scRNA-seq clusters C0 and C3), have high rates of phagocytosis and ROS production, and can directly induce cytokine production in activated T cells (Eruslanov et al., 2014), which our data suggests could occur through either CD80 or IL-23 expressed by the neutrophils. Third, we identified a gene signature suggestive of pan T17 polarization in response to dysbiosis in mice that was predictive of poor prognosis in human LUAD. T17 cells contribute to inflammation in NSCLC patients and are a poor prognostic factor (Chen et al., 2010; Jin et al., 2019a). T_H_17 and yδ T17 cells can be isolated from the blood of patients and interestingly tumor resection reduced the number of circulating cells (Bao et al., 2016), suggesting that an intratumoral source is responsible for inducing polarization. Together, these factors demonstrate that a proinflammatory tumor microenvironment in patients is reflective of the dysbiosis-associated changes we observe in KP mice.

Our results demonstrate that dysbiosis of the lung microbiome is a contributing factor in tumor development and progression, by promoting large inflammation of the microenvironment, with many known proinflammatory cytokines sharply upregulated in response to *A. temperans*. Further studies are needed with in-depth phenotypic profiling of neutrophil and T17 function in both human and mouse to precisely determine their roles in development of LUAD. Mechanistic understanding of these pathways suggests that anti-neutrophil and IL-17A therapies represent intriguing and promising targets for intervention and development of targeted therapies in LUAD.

## ACKNOWLEDGEMENTS

We thank Dr. Maria Hernandez and other staff of the NCI/CCR Single Cell Analysis Facility (funded by contract 75N91019D00024) for support with scRNA-Seq experiments. Single cell RNA sequencing and initial data analysis were conducted at the Frederick National Laboratory for Cancer Research (FNLCR) at the NCI-CCR Sequencing Facility Frederick. This research was supported in part by the NIH Intramural Research Program of NCI, a NCI/CCR FLEX Synergy award (CCH-15) to G.T. and C.C.H., and by the Israel Science Foundation, grant number 1178/20. We also thank Drs. Amanda Craig and Lichun Ma for helpful discussions and data interpretation.

## AUTHORS’ CONTRIBUTIONS

**J.K.S**: Conceptualization, data curation, formal analysis, investigation, methodology, software, visualization, writing – original draft, writing – review & editing. **N.v.M:** investigation, visualization. **C.Z:** investigation, validation. **A.I.R:** investigation, methodology. **E.V-V:** investigation, resources. **A.M:** investigation. **M.M:** investigation. **K.L.G:** conceptualization, funding acquisition, methodology. **T.C:** funding acquisition, investigation, methodology. **G.T:** conceptualization, funding acquisition, methodology, resources. **C.C.H:** conceptualization, funding acquisition, methodology, project administration, resources, supervision, writing – original draft, writing – review & editing.

## DECLARATION OF INTERESTS

The authors have nothing to disclose.

## MATERIALS AND METHODS

### Animal model, bacterial culture, and instillation

*Kras^LSL-G12D/+^; Trp53^LSL-R172H/+^* (KP) mice were purchased from Jackson Laboratory (Bar Harbor, ME) and housed under SPF conditions in accordance with the approved NIH-NCI/CCR animal use protocol (# ASP-19-334) and biosafety protocol (#19-51). Male and female KP mice aged 5-9 weeks old were administrated with 5×10^6^ PFU of Adeno-Cre-CMV virus (Viral Vector Core, University of Iowa) by intranasal instillation. *Acidovorax temperans* ATCC 49666 was obtained from the ATCC (Manassas, VA) and cultured in Nutrient Broth (BD Biosciences) at 30°C with 200 rpm shaking. Two weeks post Ad-cre instillation, mice then received six biweekly intranasal instillations of Sham (1X PBS) or *A. temperans* (1 × 10^9^ CFU). Culture inoculum was verified by serial dilution and plating on Nutrient Agar for 48 hours.

### Magnetic resonance imaging

Tumor development was determined 13 weeks post Ad-cre instillation by magnetic resonance imaging (MRI) with a 3.0T clinical scanner (Philips Intera Achieva, Best, The Netherlands). Mice were anesthetized then individually imaged using a 40-mm diameter solenoid volume receiver coil (Philips Research, Hamburg, Germany), with anesthesia and air temperature maintained at 1.5 - 2.0% isoflurane and 34-37°C, respectively. Multislice T2 weighted turbo spin echo sequence was applied in coronal view with respiratory triggering to minimize motion artifacts. The whole mouse lung was covered by an imaging slab with dimensions 38×28×16 mm. The images were acquired with a repetition time of 5333 ms, echo time of 45 ms, in plane resolution of 0.188×0.188 mm^2^, and a slice thickness 0.5mm. Lung tumor burden was analyzed by manually segmentation and volume calculation of MRI results using ITK-SNAP 3.8.0 (Krupnick et al., 2012; Yushkevich et al., 2006).

### Tissue dissociation and flow cytometry

Mice were sacrificed at 14 weeks post Ad-cre and were perfused with PBS + 2mM EDTA. Lungs were then collected, dissected, and digested at 37°C for 40 min using the Mouse Tumor Dissociation kit (Miltenyi Biotec), according to the manufacturer’s instructions. Digested lung tissues were filtered with 40 μM cell strainer to obtain a single-cell suspension. Red blood cells were lysed using RBC lysis buffer (BioLegend), according to the manufacturer’s instruction. Single cell suspensions were cryopreserved using 10% DMSO in FBS. Single cell suspensions were later thawed and stained with PE-conjugated anti-CD45 antibody (BD, clone 104, 1:100) and DAPI, then sorted on a BD FACSAria flow sorter and stored on ice.

### Single cell RNA sequencing

Approximately 7,000 cells per sample were targeted for droplet capture by 10X Chromium 3’ Dual Index v3.2 kit. Capture, cDNA synthesis, and library preparation were performed according to the manufacturer’s instructions. Sequencing was performed on Illumina NovaSeq S2 with 10 bp indices i5 and i7, 28 bp R1, and 90 bp R2 length reads. Samples were sequenced to a target depth of 50,000 reads per cell. Basecalling was performed using RTA v2.4.11, demultiplexing with Bcl2fastq v2.20, and read alignment to mouse genome version mm10, tagging, gene and transcript counting, and clustering analysis were performed using CellRanger v6.0.2. The generated filtered matrices were used for downstream analysis.

### Quality control and single cell data analysis

Cells with less than 100 features, more than 2800 features, or greater than 5% mitochondrial reads were excluded using Seurat v4.1.0 (Hao et al., 2021), followed by removal of fibroblasts (n = 590), leaving a total 25,477 cells with 20,737 genes detected in at least one cell. Average features per cell was 1,397 with an average UMI of 3,788 per cell. Passing cell counts were normalized and scaled, then neighbors were clustered using the first 50 dimensions with a resolution of 0.8, resulting in a total of 22 clusters. Cell types were manually assigned using standard marker genes and the ImmGen database (Heng et al., 2008). Both the slingshot (v2.2.1) and monocle2 (v2.22.0) packages were used for trajectory and pseudotime analysis (Qiu et al., 2017; Street et al., 2018). Differentially expressed genes were identified using the Seurat FindMarkers() function and then ssGSEA was performed using the escape v1.4.0 (Borcherding et al., 2021). Cell-cell communication was predicted using CellChat v1.1.2 (Jin et al., 2021), with the Cell-Cell Interactions and Secreted Signaling datasets selected.

### Survival analysis

Gene symbols for the murine T17 signature were converted to their human orthologues using biomaRt v2.50.3 (Durinck et al., 2009). The resulting gene lists were subsetted from expression matrices for GSE30210, GSE31210, GSE50081, and TCGA (Collisson et al., 2014; Der et al., 2014; Okayama et al., 2012; Rousseaux et al., 2013). Human T17 signature scores were tested in a Cox analysis and then cutoff values for low and high expression in each cohort were separately determined using maximally selected rank statistics within survminer v0.4.9 (https://cran.r-project.org/web/packages/survminer/index.html).

### Statistical analysis

Statistical analyses of lung tumor burden and lung weight were performed using GraphPad Prism 8. All sequencing analyses were performed in R v4.1.0 (http://www.r-project.org/). Unless otherwise stated, the Mann-Whitney test was used for comparison between two groups and the Kruskal-Wallis test was used for comparison between three or more groups. P < 0.05 was considered as significantly different.

### Availability of data and materials

Single-cell RNA-sequencing data generated in this study were deposited in the NCBI GEO database under the accession number GEO: GSE207477. Scripts used in analysis are available upon request from the corresponding author.

## SUPPLEMENTAL FIGURE AND TABLE LEGENDS

**Fig. S1 – MHC I is broadly expressed in MoMaDCs and upregulated in TAMs.**

(A) Dotplot of M1 and M2 macrophage marker gene expression in MoMaDCs.

(B) Dotplot of individual MHC class I and II gene expression in MoMaDCs.

(C) Comparison of average expression of each MHC I component gene (*H2-D1, -K1, -M2, -M3, -Q4, -Q6, -Q7, -T22, -T23*) by treatment for each cell type.

Data presented as median value plus quartiles for boxplots, n.s. not significant, * *p* < 0.05, **** *p* < 0.0001.

**Fig. S2 – *A. temperans* induces MHC II upregulation in alveolar macrophages.**

(A) UMAP plots of alveolar macrophages (AMs) cell types (left) and treatment groups (right).

(B) Barplot of the relative abundance for each AM cluster by treatment group.

(C) Dotplot of marker genes for each cluster.

(D) Density plot for AM marker genes (*Mrc1* and *Tgm2*) and monocyte-derived AMs (*Msr1* and *Ccr2*).

(E) Quantification of the total expression of marker genes by treatment.

(F) ssGSEA heatmap of average normalized enrichment scores for each AM cluster.

(G) Dotplot of M1 and M2 macrophage marker gene expression in AMs.

(H) Dotplot of individual MHC class I and II gene expression in AMs.

(I) Comparison of average expression of each MHC I component gene (*H2-D1, -K1, -M2, -M3, - Q4, -Q6, -Q7, -T22, -T23*) by treatment for each cluster.

(J) Comparison of average expression of each MHC II component gene (*H2-Aa, -Ab1, -DMa, - DMb1, -DMb2, -Eb1, -Eb2, -Oa, -Ob*) by treatment for each cell type. Data presented as median value plus quartiles for boxplots, n.s. not significant, * *p* < 0.05, **** *p* < 0.0001.

**Fig. S3 – Known DN T cell subtypes are not detected.**

(A) Density plots for ILC (left), MAIT (center), and NKT (right) cell markers.

(B) UMAP plots of signature scores for ILC (left), MAIT (center), and NKT cells (right).

**Fig. S4 – Bacterial infection datasets reveal *A. temperans* induces specific T_H_17 and general IFN-γ response in T cells.**

(A) UMAP plot of a T_H_17 response gene signature in a *Citrobacter rodentium* infection model [44] within T cell subtypes.

(B) Violin plot of the expression of the *C. rodentium* gene signature by T cell subtype.

(C) UMAP plot of an IFN-γ response gene signature in a *Salmonella enteria* serovar Typhimurium infection model [44] within T cell subtypes.

(D) Violin plot of the expression of the *Salmonella* Typhimurium gene signature by T cell subtype.

**Fig. S5 – Both IFN-γ and IL-17 are expressed in response to *A. temperans*.**

(A) Density plot of the cytokine *Il17a* and its transcription factor *Rorc* (left) and the cytokine *Ifng* and its transcription factor *Stat4* (right) within T cells.

(B) Expression level of tissue residency marker genes by T cell subtype.

(C) Percent of cells *Ifng* positive per T cell subtype and treatment group.

(D and E) Expression level of *Il17a* (right) and *Ifng* (left) within the Th17 (D) and γδ T (E) clusters by treatment group.

(F) Correlation of *Il17a* and *Ifng* in all T cells.

(G) Barplot of *Il17a/Ifng* expression in SP, DP, and DN T cells by treatment.

**Fig. S6 – Specificity of ligand-receptor signaling pathways by treatment group.**

Relative (left) and absolute (right) contribution of aggregate signaling pathways, ordered from sham-exclusive (top, blue) to *A. temperans*-exclusive (bottom, pink).

**Table S1 – Main cell type marker genes.**

**Table S2 – MoMaDC marker genes.**

**Table S3 – Alveolar macrophage marker genes.**

**Table S4 – Neutrophil marker genes.**

**Table S5 – T cell marker genes.**

**Table S6 – Common Th17 and γδ T marker genes used for pan T17 signature.**

**Table S7 – Cell-cell interactions increased in *A. temperans* mice.**

**Table S8 – Cell-cell interactions decreased in *A. temperans* mice.**

## Notes

https://www.ncbi.nlm.nih.gov/geo/query/acc.cgi?acc=GSE207477

### Competing Interest Statement

The authors have declared no competing interest.

## References

Abbas, A., Vu Manh, T.-P., Valente, M., Collinet, N., Attaf, N., Dong, C., Naciri, K., Chelbi, R., Brelurut, G., Cervera-Marzal, I., et al. (2020). The activation trajectory of plasmacytoid dendritic cells in vivo during a viral infection. Nature Immunology 21, 983–997.

Alshetaiwi, H., Pervolarakis, N., McIntyre, L.L., Ma, D., Nguyen, Q., Rath, J.A., Nee, K., Hernandez, G., Evans, K., Torosian, L., et al. (2020). Defining the emergence of myeloid-derived suppressor cells in breast cancer using single-cell transcriptomics. Science Immunology 5, eaay6017.

Ancey, P.-B., Contat, C., Boivin, G., Sabatino, S., Pascual, J., Zangger, N., Perentes, J.Y., Peters, S., Abel, E.D., Kirsch, D.G., et al. (2021). GLUT1 expression in tumor-associated neutrophils promotes lung cancer growth and resistance to radiotherapy. Cancer Research 81, 2345–2357.

Bao, Z., Lu, G., Cui, D., Yao, Y., Yang, G., and Zhou, J. (2016). IL-17A-producing T cells are associated with the progression of lung adenocarcinoma. Oncol Rep 36, 641–650.

Borcherding, N., Vishwakarma, A., Voigt, A.P., Bellizzi, A., Kaplan, J., Nepple, K., Salem, A.K., Jenkins, R.W., Zakharia, Y., and Zhang, W. (2021). Mapping the immune environment in clear cell renal carcinoma by single-cell genomics. Communications Biology 4, 122.

Bracke, K.R., D’hulst, A.I., Maes, T., Moerloose, K.B., Demedts, I.K., Lebecque, S., Joos, G.F., and Brusselle, G.G. (2006). Cigarette Smoke-Induced Pulmonary Inflammation and Emphysema Are Attenuated in CCR6-Deficient Mice. 177, 4350–4359.

Bray, F., Ferlay, J., Soerjomataram, I., Siegel, R.L., Torre, L.A., and Jemal, A. (2018). Global cancer statistics 2018: GLOBOCAN estimates of incidence and mortality worldwide for 36 cancers in 185 countries. CA Cancer J Clin 68, 394–424.

Brenner, A.V., Wang, Z., Kleinerman, R.A., Wang, L., Zhang, S., Metayer, C., Chen, K., Lei, S., Cui, H., and Lubin, J.H. (2001). Previous pulmonary diseases and risk of lung cancer in Gansu Province, China. International Journal of Epidemiology 30, 118–124.

Cavalcanti, D.M.H., Lotufo, C.M.C., Borelli, P., Ferreira, Z.S., Markus, R.P., and Farsky, S.H.P. (2007). Endogenous glucocorticoids control neutrophil mobilization from bone marrow to blood and tissues in non-inflammatory conditions. British Journal of Pharmacology 152, 1291–1300.

Chen, X., Wan, J., Liu, J., Xie, W., Diao, X., Xu, J., Zhu, B., and Chen, Z. (2010). Increased IL-17-producing cells correlate with poor survival and lymphangiogenesis in NSCLC patients. Lung Cancer 69, 348–354.

Ciofani, M., Madar, A., Galan, C., Sellars, M., Mace, K., Pauli, F., Agarwal, A., Huang, W., Parkurst, Christopher N., Muratet, M., et al. (2012). A validated regulatory network for Th17 cell specification. Cell 151, 289–303.

Coffelt, S.B., Wellenstein, M.D., and de Visser, K.E. (2016). Neutrophils in cancer: neutral no more. Nature Reviews Cancer 16, 431–446.

Collisson, E.A., Campbell, J.D., Brooks, A.N., Berger, A.H., Lee, W., Chmielecki, J., Beer, D.G., Cope, L., Creighton, C.J., Danilova, L., et al. (2014). Comprehensive molecular profiling of lung adenocarcinoma. Nature 511, 543–550.

Cortez-Retamozo, V., Etzrodt, M., Newton, A., Rauch, P.J., Chudnovskiy, A., Berger, C., Ryan, R.J.H., Iwamoto, Y., Marinelli, B., Gorbatov, R., et al. (2012). Origins of tumor-associated macrophages and neutrophils. Proceedings of the National Academy of Sciences 109, 2491–2496.

Der, S.D., Sykes, J., Pintilie, M., Zhu, C.-Q., Strumpf, D., Liu, N., Jurisica, I., Shepherd, F.A., and Tsao, M.-S. (2014). Validation of a histology-independent prognostic gene signature for early-stage, non–small-cell lung cancer including stage IA patients. Journal of Thoracic Oncology 9, 59–64.

Dickson, R.P., Erb-Downward, J.R., Freeman, C.M., McCloskey, L., Falkowski, N.R., Huffnagle, G.B., and Curtis, J.L. (2017). Bacterial topography of the healthy human lower respiratory tract. mBio 8, e02287–02216.

Durinck, S., Spellman, P.T., Birney, E., and Huber, W. (2009). Mapping identifiers for the integration of genomic datasets with the R/Bioconductor package biomaRt. Nature Protocols 4, 1184–1191.

Engblom, C., Pfirschke, C., Zilionis, R., Da Silva Martins, J., Bos, S.A., Courties, G., Rickelt, S., Severe, N., Baryawno, N., Faget, J., et al. (2017). Osteoblasts remotely supply lung tumors with cancer-promoting SiglecFhigh neutrophils. Science 358, eaal5081.

Engels, E.A., Shen, M., Chapman, R.S., Pfeiffer, R.M., Yu, Y.-Y., He, X., and Lan, Q. (2009). Tuberculosis and subsequent risk of lung cancer in Xuanwei, China. 124, 1183–1187.

Eruslanov, E.B., Bhojnagarwala, P.S., Quatromoni, J.G., Stephen, T.L., Ranganathan, A., Deshpande, C., Akimova, T., Vachani, A., Litzky, L., Hancock, W.W., et al. (2014). Tumor-associated neutrophils stimulate T cell responses in early-stage human lung cancer. The Journal of Clinical Investigation 124, 5466–5480.

Faget, J., Groeneveld, S., Boivin, G., Sankar, M., Zangger, N., Garcia, M., Guex, N., Zlobec, I., Steiner, L., Piersigilli, A., et al. (2017). Neutrophils and snail orchestrate the establishment of a pro-tumor microenvironment in lung cancer. Cell Reports 21, 3190–3204.

Giese, M.A., Hind, L.E., and Huttenlocher, A. (2019). Neutrophil plasticity in the tumor microenvironment. Blood 133, 2159–2167.

Greathouse, K.L., White, J.R., Vargas, A.J., Bliskovsky, V.V., Beck, J.A., von Muhlinen, N., Polley, E.C., Bowman, E.D., Khan, M.A., Robles, A.I., et al. (2018). Interaction between the microbiome and TP53 in human lung cancer. Genome Biol 19.

Hao, Y., Hao, S., Andersen-Nissen, E., Mauck, W.M., III, Zheng, S., Butler, A., Lee, M.J., Wilk, A.J., Darby, C., Zager, M., et al. (2021). Integrated analysis of multimodal single-cell data. Cell 184, 3573–3587.e3529.

Hasday, J.D., Bascom, R., Costa, J.J., Fitzgerald, T., and Dubin, W. (1999). Bacterial Endotoxin Is an Active Component of Cigarette Smoke. CHEST 115, 829–835.

Hecht, S.S. (2012). Lung carcinogenesis by tobacco smoke. 131, 2724–2732.

Hedrick, C.C., and Malanchi, I. (2022). Neutrophils in cancer: heterogeneous and multifaceted. Nature Reviews Immunology 22, 173–187.

Heng, T.S.P., Painter, M.W., Elpek, K., Lukacs-Kornek, V., Mauermann, N., Turley, S.J., Koller, D., Kim, F.S., Wagers, A.J., Asinovski, N., et al. (2008). The Immunological Genome Project: networks of gene expression in immune cells. Nature Immunology 9, 1091–1094.

Horton, B.L., Morgan, D.M., Momin, N., Zagorulya, M., Torres-Mejia, E., Bhandarkar, V., Wittrup, K.D., Love, J.C., and Spranger, S. (2021). Lack of CD8+ T cell effector differentiation during priming mediates checkpoint blockade resistance in non-small cell lung cancer. Science Immunology 6, eabi8800.

Huang, C., and Shi, G. (2019). Smoking and microbiome in oral, airway, gut and some systemic diseases. Journal of Translational Medicine 17, 225.

Jackson, E.L., Olive, K.P., Tuveson, D.A., Bronson, R., Crowley, D., Brown, M., and Jacks, T. (2005). The Differential Effects of Mutant p53 Alleles on Advanced Murine Lung Cancer. 65, 10280–10288.

Jackson, E.L., Willis, N., Mercer, K., Bronson, R.T., Crowley, D., Montoya, R., Jacks, T., and Tuveson, D.A. (2001). Analysis of lung tumor initiation and progression using conditional expression of oncogenic K-ras. Gene Dev 15, 3243–3248.

Jin, C., Lagoudas, G.K., Zhao, C., Bullman, S., Bhutkar, A., Hu, B., Ameh, S., Sandel, D., Liang, X.S., Mazzilli, S., et al. (2019a). Commensal Microbiota Promote Lung Cancer Development via gammadelta T Cells. Cell 176, 998–1013 e1016.

Jin, J., Gan, Y., Liu, H., Wang, Z., Yuan, J., Deng, T., Zhou, Y., Zhu, Y., Zhu, H., Yang, S., et al. (2019b). Diminishing microbiome richness and distinction in the lower respiratory tract of lung cancer patients: A multiple comparative study design with independent validation. Lung Cancer 136, 129–135.

Jin, S., Guerrero-Juarez, C.F., Zhang, L., Chang, I., Ramos, R., Kuan, C.-H., Myung, P., Plikus, M.V., and Nie, Q. (2021). Inference and analysis of cell-cell communication using CellChat. Nature Communications 12, 1088.

Kargl, J., Busch, S.E., Yang, G.H.Y., Kim, K.-H., Hanke, M.L., Metz, H.E., Hubbard, J.J., Lee, S.M., Madtes, D.K., McIntosh, M.W., et al. (2017). Neutrophils dominate the immune cell composition in non-small cell lung cancer. Nature Communications 8, 14381.

Kim, M.-H., Granick, J.L., Kwok, C., Walker, N.J., Borjesson, D.L., Curry, F.-R.E., Miller, L.S., and Simon, S.I. (2011). Neutrophil survival and c-kit+-progenitor proliferation in Staphylococcus aureus–infected skin wounds promote resolution. Blood 117, 3343–3352.

Kiner, E., Willie, E., Vijaykumar, B., Chowdhary, K., Schmutz, H., Chandler, J., Schnell, A., Thakore, P.I., LeGros, G., Mostafavi, S., et al. (2021). Gut CD4+ T cell phenotypes are a continuum molded by microbes, not by TH archetypes. Nature Immunology 22, 216–228.

Kroon, E.E., Coussens, A.K., Kinnear, C., Orlova, M., Möller, M., Seeger, A., Wilkinson, R.J., Hoal, E.G., and Schurr, E. (2018). Neutrophils: Innate Effectors of TB Resistance? Frontiers in Immunology 9.

Krupnick, A.S., Tidwell, V.K., Engelbach, J.A., Alli, V.V., Nehorai, A., You, M., Vikis, H.G., Gelman, A.E., Kreisel, D., and Garbow, J.R. (2012). Quantitative monitoring of mouse lung tumors by magnetic resonance imaging. Nat Protoc 7, 128–142.

Künzli, M., Schreiner, D., Pereboom, T.C., Swarnalekha, N., Litzler, L.C., Lötscher, J., Ertuna, Y.I., Roux, J., Geier, F., Jakob, R.P., et al. (2020). Long-lived T follicular helper cells retain plasticity and help sustain humoral immunity. Science Immunology 5, eaay5552.

Larsson, L., Szponar, B., Ridha, B., Pehrson, C., Dutkiewicz, J., Krysińska-Traczyk, E., and Sitkowska, J. (2008). Identification of bacterial and fungal components in tobacco and tobacco smoke. Tobacco Induced Diseases 4, 4.

Leng, Q., Holden, V.K., Deepak, J., Todd, N.W., and Jiang, F. (2021). Microbiota Biomarkers for Lung Cancer. Diagnostics 11, 407.

Maier, B., Leader, A.M., Chen, S.T., Tung, N., Chang, C., LeBerichel, J., Chudnovskiy, A., Maskey, S., Walker, L., Finnigan, J.P., et al. (2020). A conserved dendritic-cell regulatory program limits antitumour immunity. Nature 580, 257–262.

Malayil, L., Chattopadhyay, S., Kulkarni, P., Hittle, L., Clark, P.I., Mongodin, E.F., and Sapkota, A.R. (2020). Mentholation triggers brand-specific shifts in the bacterial microbiota of commercial cigarette products. Applied Microbiology and Biotechnology 104, 6287–6297.

Melkamu, T., Qian, X., Upadhyaya, P., O'Sullivan, M.G., and Kassie, F. (2013). Lipopolysaccharide Enhances Mouse Lung Tumorigenesis:A Model for Inflammation-Driven Lung Cancer. Veterinary Pathology 50, 895–902.

Minoda, Y., Virshup, I., Leal Rojas, I., Haigh, O., Wong, Y., Miles, J.J., Wells, C.A., and Radford, K.J. (2017). Human CD141+ dendritic cell and CD1c+ dendritic cell undergo concordant early genetic programming after activation in humanized mice In vivo. Frontiers in Immunology 8, 1419.

Miragaia, R.J., Gomes, T., Chomka, A., Jardine, L., Riedel, A., Hegazy, A.N., Whibley, N., Tucci, A., Chen, X., Lindeman, I., et al. (2019). Single-cell transcriptomics of regulatory T cells reveals trajectories of tissue adaptation. Immunity 50, 493–504.e497.

Mollaoglu, G., Jones, A., Wait, S.J., Mukhopadhyay, A., Jeong, S., Arya, R., Camolotto, S.A., Mosbruger, T.L., Stubben, C.J., Conley, C.J., et al. (2018). The lineage-defining transcription factors SOX2 and NKX2-1 determine lung cancer cell fate and shape the tumor immune microenvironment. Immunity 49, 764–779.e769.

O'Callaghan, D.S., O'Donnell, D., O'Connell, F., and O'Byrne, K.J. (2010). The Role of Inflammation in the Pathogenesis of Non-small Cell Lung Cancer. Journal of Thoracic Oncology 5, 2024–2036.

Okayama, H., Kohno, T., Ishii, Y., Shimada, Y., Shiraishi, K., Iwakawa, R., Furuta, K., Tsuta, K., Shibata, T., Yamamoto, S., et al. (2012). Identification of genes upregulated in ALK-positive and EGFR/KRAS/ALK-negative lung adenocarcinomas. Cancer Research 72, 100–111.

Pfirschke, C., Engblom, C., Gungabeesoon, J., Lin, Y., Rickelt, S., Zilionis, R., Messemaker, M., Siwicki, M., Gerhard, G.M., Kohl, A., et al. (2020). Tumor-Promoting Ly-6G+ SiglecFhigh Cells Are Mature and Long-Lived Neutrophils. Cell Reports 32.

Qiu, X., Mao, Q., Tang, Y., Wang, L., Chawla, R., Pliner, H.A., and Trapnell, C. (2017). Reversed graph embedding resolves complex single-cell trajectories. Nature Methods 14, 979–982.

Ramírez-Labrada, A.G., Isla, D., Artal, A., Arias, M., Rezusta, A., Pardo, J., and Gálvez, E.M. (2020). The Influence of Lung Microbiota on Lung Carcinogenesis, Immunity, and Immunotherapy. Trends in Cancer 6, 86–97.

Rapoport, B.L., Theron, A.J., Vorobiof, Daniel A., Langenhoven, L., Hall, J.M., Van Eeden, R.I., Smit, T., Chan, S.-W., Botha, M.C., Ratts, J.I., et al. (2020). Prognostic significance of the neutrophil/lymphocyte ratio in patients undergoing treatment with nivolumab for recurrent non-small-cell lung cancer. Lung Cancer Management 9, LMT37.

Rousseaux, S., Debernardi, A., Jacquiau, B., Vitte, A.-L., Vesin, A., Nagy-Mignotte, H., Moro-Sibilot, D., Brichon, P.-Y., Lantuejoul, S., Hainaut, P., et al. (2013). Ectopic activation of germline and placental genes identifies aggressive metastasis-prone lung cancers. Science Translational Medicine 5, 186ra166–186ra166.

Segal, L.N., Clemente, J.C., Tsay, J.-C.J., Koralov, S.B., Keller, B.C., Wu, B.G., Li, Y., Shen, N., Ghedin, E., Morris, A., et al. (2016). Enrichment of the lung microbiome with oral taxa is associated with lung inflammation of a Th17 phenotype. Nature Microbiology 1, 16031.

Shiels, M.S., Albanes, D., Virtamo, J., and Engels, E.A. (2011). Increased Risk of Lung Cancer in Men with Tuberculosis in the Alpha-Tocopherol, Beta-Carotene Cancer Prevention Study. 20, 672–678.

Shimizu, M., Miyanaga, A., Seike, M., Matsuda, K., Matsumoto, M., Noro, R., Fujita, K., Mano, Y., Furuya, N., Kubota, K., et al. (2022). The respiratory microbiome associated with chronic obstructive pulmonary disease comorbidity in non-small cell lung cancer. Thoracic Cancer n/a, 1–8.

Siegel, R.L., Miller, K.D., and Jemal, A. (2018). Cancer statistics, 2018. CA Cancer J Clin 68, 7–30.

Simoncello, F., Piperno, G.M., Caronni, N., Amadio, R., Cappelletto, A., Canarutto, G., Piazza, S., Bicciato, S., and Benvenuti, F. (2022). CXCL5-mediated accumulation of mature neutrophils in lung cancer tissues impairs the differentiation program of anticancer CD8 T cells and limits the efficacy of checkpoint inhibitors. OncoImmunology 11, 2059876.

Singleton, D.R., Guzmán Ramirez, L., and Aitken, M.D. (2009). Characterization of a polycyclic aromatic hydrocarbon degradation gene cluster in a phenanthrene-degrading Acidovorax strain. Applied and Environmental Microbiology 75, 2613–2620.

Street, K., Risso, D., Fletcher, R.B., Das, D., Ngai, J., Yosef, N., Purdom, E., and Dudoit, S. (2018). Slingshot: cell lineage and pseudotime inference for single-cell transcriptomics. BMC Genomics 19, 477.

Tsay, J.-C.J., Wu, B.G., Sulaiman, I., Gershner, K., Schluger, R., Li, Y., Yie, T.-A., Meyn, P., Olsen, E., Perez, L., et al. (2021). Lower Airway Dysbiosis Affects Lung Cancer Progression. Cancer Discovery 11, 293–307.

Woodfin, A., Beyrau, M., Voisin, M.-B., Ma, B., Whiteford, J.R., Hordijk, P.L., Hogg, N., and Nourshargh, S. (2016). ICAM-1-expressing neutrophils exhibit enhanced effector functions in murine models of endotoxemia. Blood 127, 898–907.

Xavier, J.B., Young, V.B., Skufca, J., Ginty, F., Testerman, T., Pearson, A.T., Macklin, P., Mitchell, A., Shmulevich, I., Xie, L., et al. (2020). The Cancer Microbiome: Distinguishing Direct and Indirect Effects Requires a Systemic View. Trends in Cancer 6, 192–204.

Xu, W., Li, R., and Sun, Y. (2019). Increased IFN-γ-producing Th17/Th1 cells and their association with lung function and current smoking status in patients with chronic obstructive pulmonary disease. BMC Pulmonary Medicine 19, 137.

Yamaguchi, T., Yanagisawa, K., Sugiyama, R., Hosono, Y., Shimada, Y., Arima, C., Kato, S., Tomida, S., Suzuki, M., Osada, H., et al. (2012). NKX2-1/TITF1/TTF-1-Induced ROR1 is required to sustain EGFR survival signaling in lung adenocarcinoma. Cancer Cell 21, 348–361.

Yu, G., Gail, M.H., Consonni, D., Carugno, M., Humphrys, M., Pesatori, A.C., Caporaso, N.E., Goedert, J.J., Ravel, J., and Landi, M.T. (2016). Characterizing human lung tissue microbiota and its relationship to epidemiological and clinical features. Genome Biol 17, 163.

Yushkevich, P.A., Piven, J., Hazlett, H.C., Smith, R.G., Ho, S., Gee, J.C., and Gerig, G. (2006). User-guided 3D active contour segmentation of anatomical structures: significantly improved efficiency and reliability. Neuroimage 31, 1116–1128.

Zhang, L., Li, Z., Skrzypczynska, K.M., Fang, Q., Zhang, W., O'Brien, S.A., He, Y., Wang, L., Zhang, Q., Kim, A., et al. (2020). Single-cell analyses inform mechanisms of myeloid-targeted therapies in colon cancer. Cell 181, 442–459.e429.

