## Supplemental Files for "*Acidovorax temperans* polarizes T17 cells and skews neutrophil maturation to promote lung adenocarcinoma development"

### Supplementary Figure 1

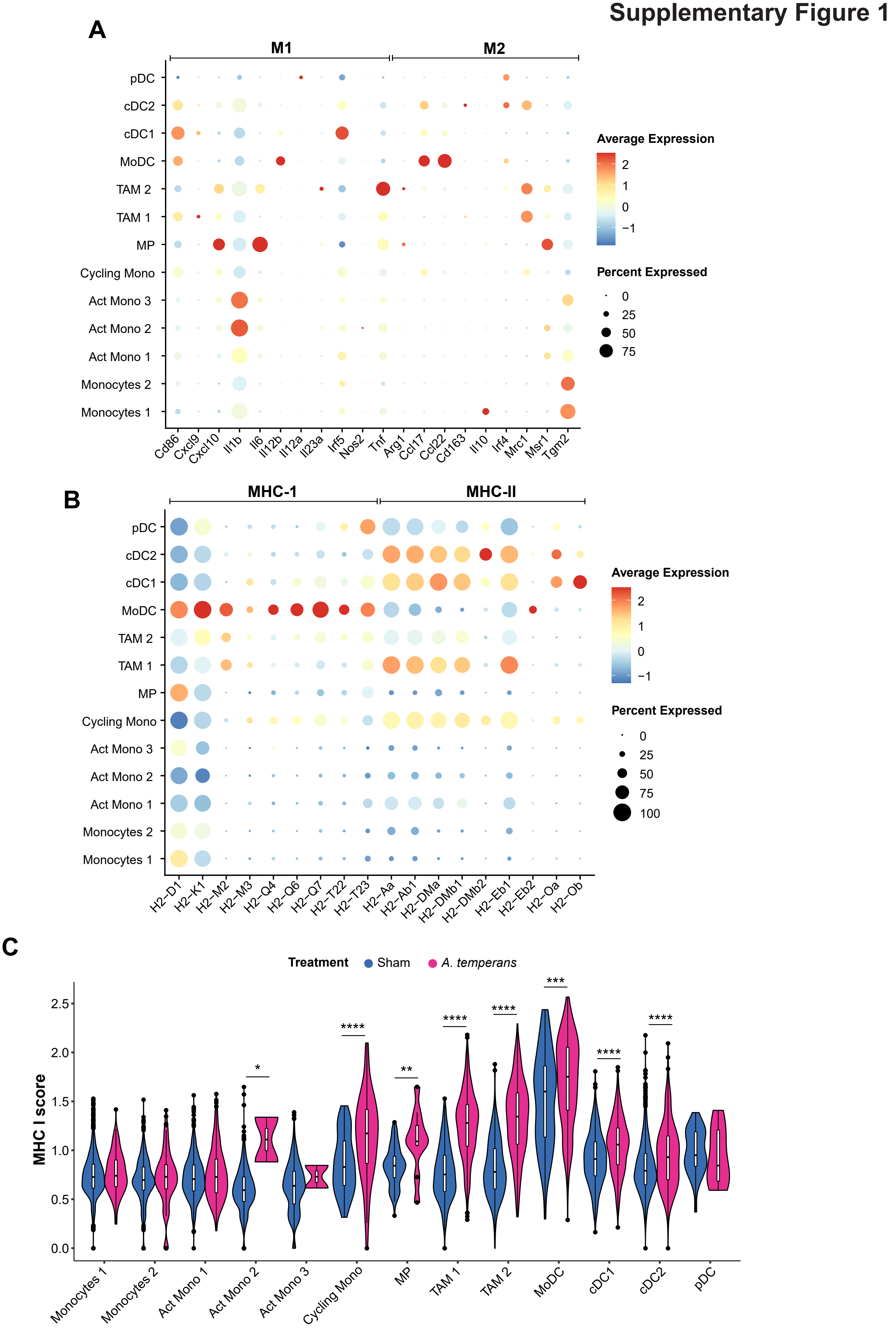

**Fig. S1 – MHC I is broadly expressed in MoMaDCs and upregulated in TAMs.**
(A) Dotplot of M1 and M2 macrophage marker gene expression in MoMaDCs.
(B) Dotplot of individual MHC class I and II gene expression in MoMaDCs.
(C) Comparison of average expression of each MHC I component gene (*H2-D1, -K1, -M2, -M3, -Q4, -Q6, -Q7, -T22, -T23*) by treatment for each cell type.
Data presented as median value plus quartiles for boxplots, n.s. not significant, * $p < 0.05$, **** $p < 0.0001$.

### Supplementary Figure 2

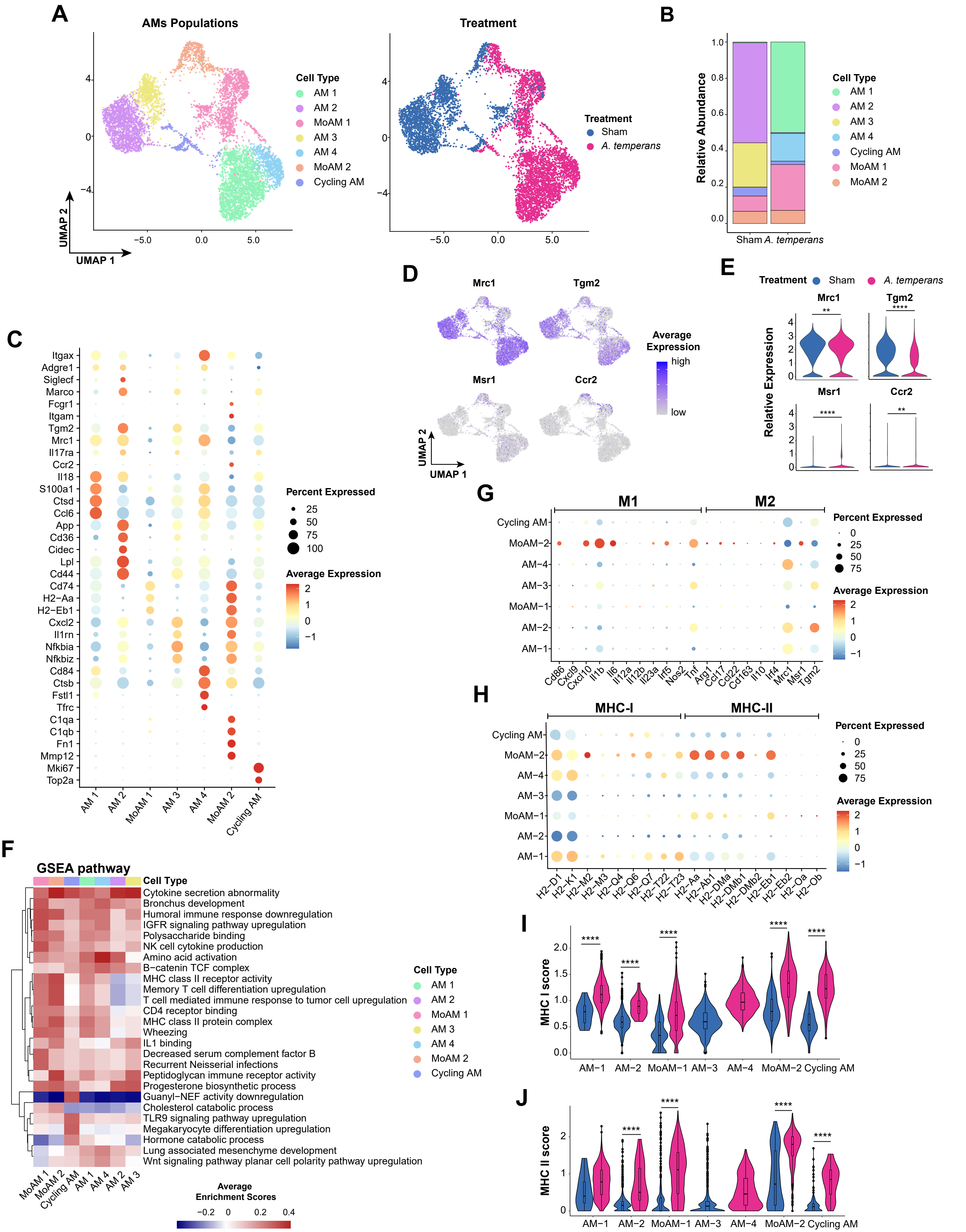

**Fig. S2 – *A. temperans* induces MHC II upregulation in alveolar macrophages.**
(A) UMAP plots of alveolar macrophages (AMs) cell types (left) and treatment groups (right).
(B) Barplot of the relative abundance for each AM cluster by treatment group.
(C) Dotplot of marker genes for each cluster.
(D) Density plot for AM marker genes (*Mrc1* and *Tgm2*) and monocyte-derived AMs (*Msr1* and *Ccr2*).
(E) Quantification of the total expression of marker genes by treatment.
(F) ssGSEA heatmap of average normalized enrichment scores for each AM cluster.
(G) Dotplot of M1 and M2 macrophage marker gene expression in AMs.
(H) Dotplot of individual MHC class I and II gene expression in AMs.
(I) Comparison of average expression of each MHC I component gene (*H2-D1, -K1, -M2, -M3, -Q4, -Q6, -Q7, -T22, -T23*) by treatment for each cluster.
(J) Comparison of average expression of each MHC II component gene (*H2-Aa, -Ab1, -DMa, -DMb1, -DMb2, -Eb1, -Eb2, -Oa, -Ob*) by treatment for each cell type.
Data presented as median value plus quartiles for boxplots, n.s. not significant, * $p < 0.05$, **** $p < 0.0001$.

### Supplementary Figure 3

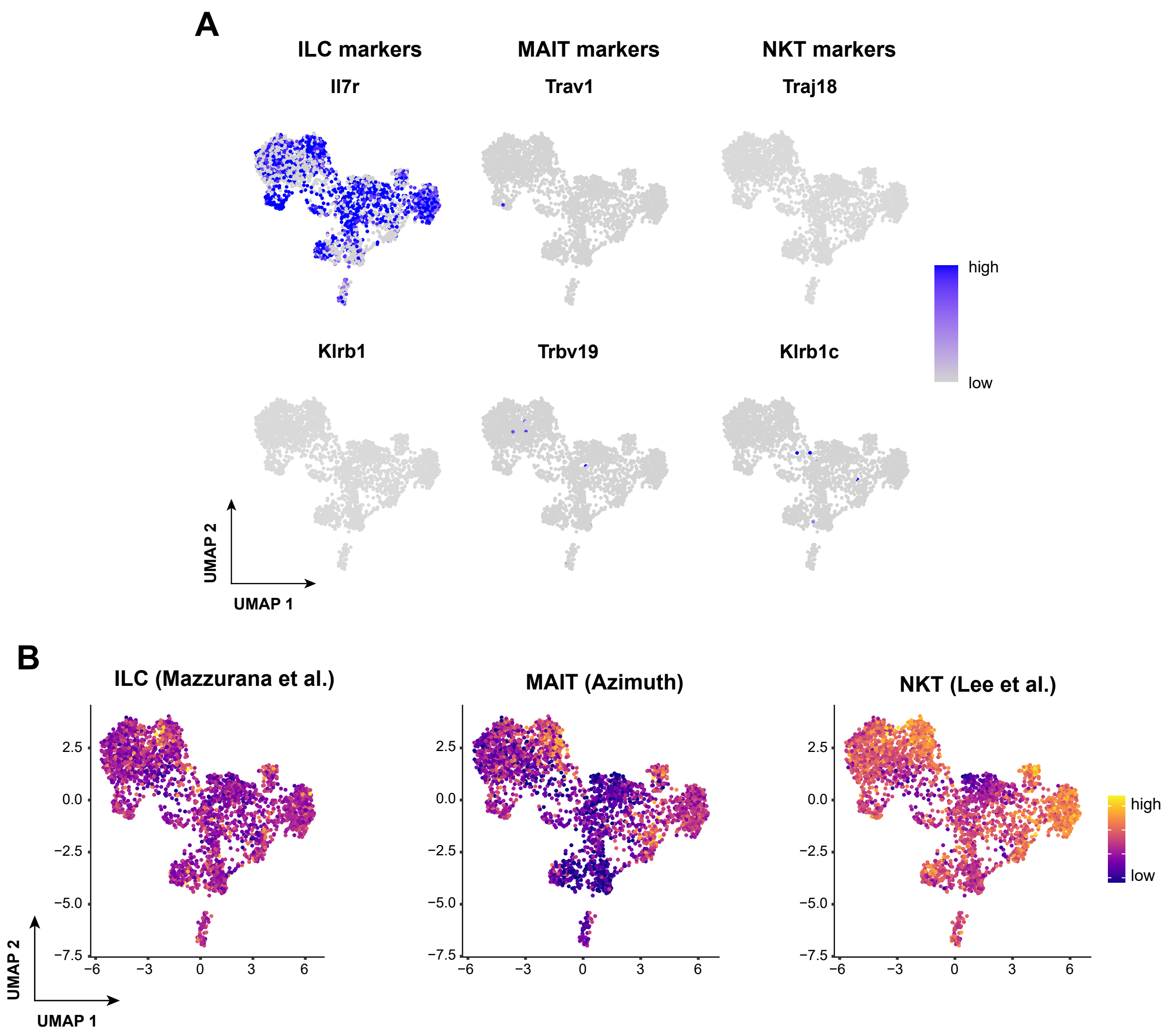

**Fig. S3 – Known DN T cell subtypes are not detected.**
(A) Density plots for ILC (left), MAIT (center), and NKT (right) cell markers.
(B) UMAP plots of signature scores for ILC (left), MAIT (center), and NKT cells (right).

#### Supplementary Figure 4

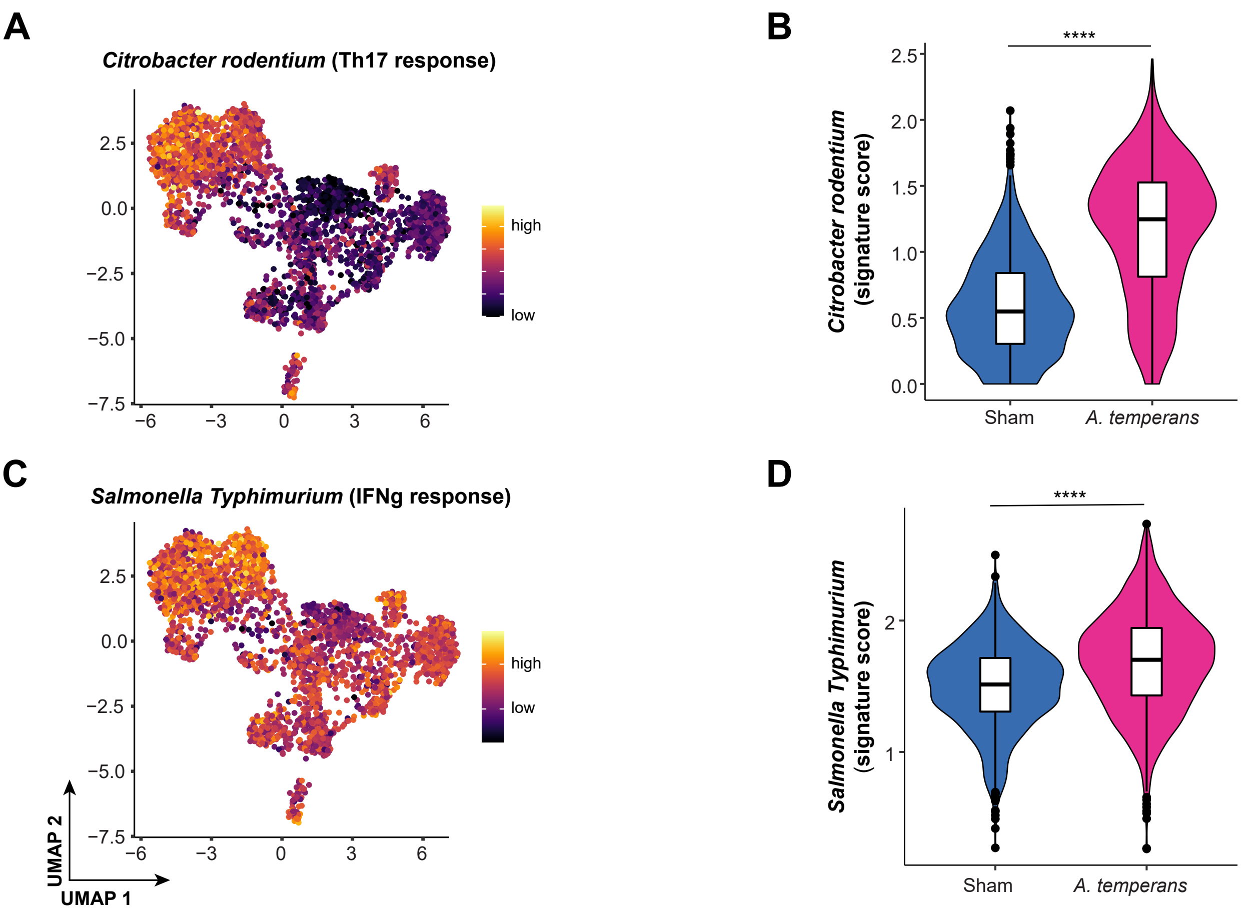

**Fig. S4 – Bacterial infection datasets reveal *A. temperans* induces specific $T_H17$ and general IFN-γ response in T cells.**

(A) UMAP plot of a $T_H17$ response gene signature in a *Citrobacter rodentium* infection model [44] within T cell subtypes.

(B) Violin plot of the expression of the *C. rodentium* gene signature by T cell subtype.

(C) UMAP plot of an IFN-γ response gene signature in a *Salmonella enteria* serovar Typhimurium infection model [44] within T cell subtypes.

(D) Violin plot of the expression of the *S.* Typhimurium gene signature by T cell subtype.

Data presented as median value plus quartiles for boxplots, **** $p < 0.0001$.

### Supplementary Figure 5

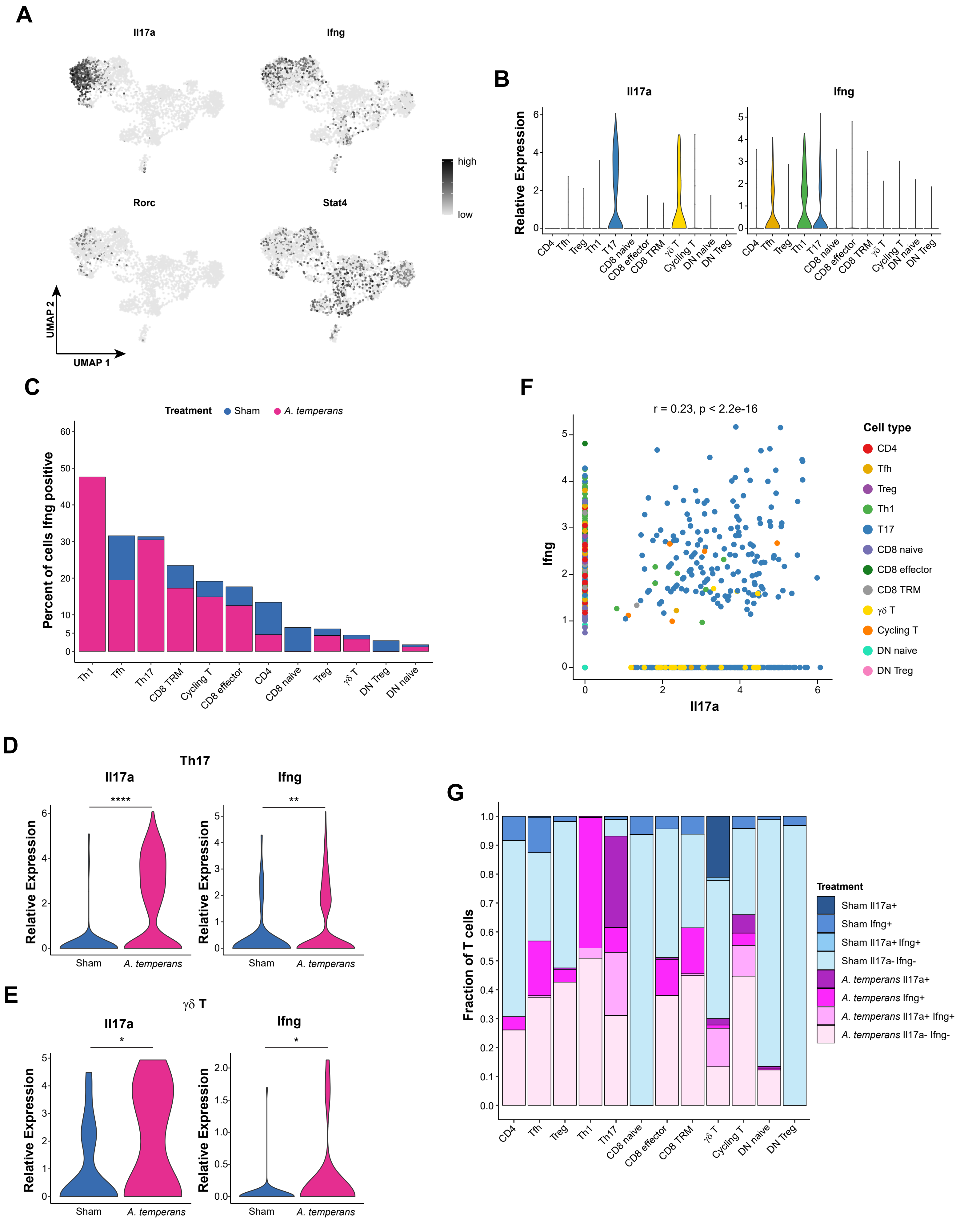

**Fig. S5 – Both IFN-γ and IL-17 are expressed in response to *A. temperans*.**
(A) Density plot of the cytokine *Il17a* and its transcription factor *Rorc* (left) and the cytokine *Ifng* and its transcription factor *Stat4* (right) within T cells.
(B) Expression level of tissue residency marker genes by T cell subtype.
(C) Percent of cells *Ifng* positive per T cell subtype and treatment group.
(D and E) Expression level of *Il17a* (right) and *Ifng* (left) within the Th17 (D) and γδ T (E) clusters by treatment group.
(F) Correlation of *Il17a* and *Ifng* in all T cells.
(G) Barplot of *Il17a/Ifng* expression in SP, DP, and DN T cells by treatment.

### Supplementary Figure 6

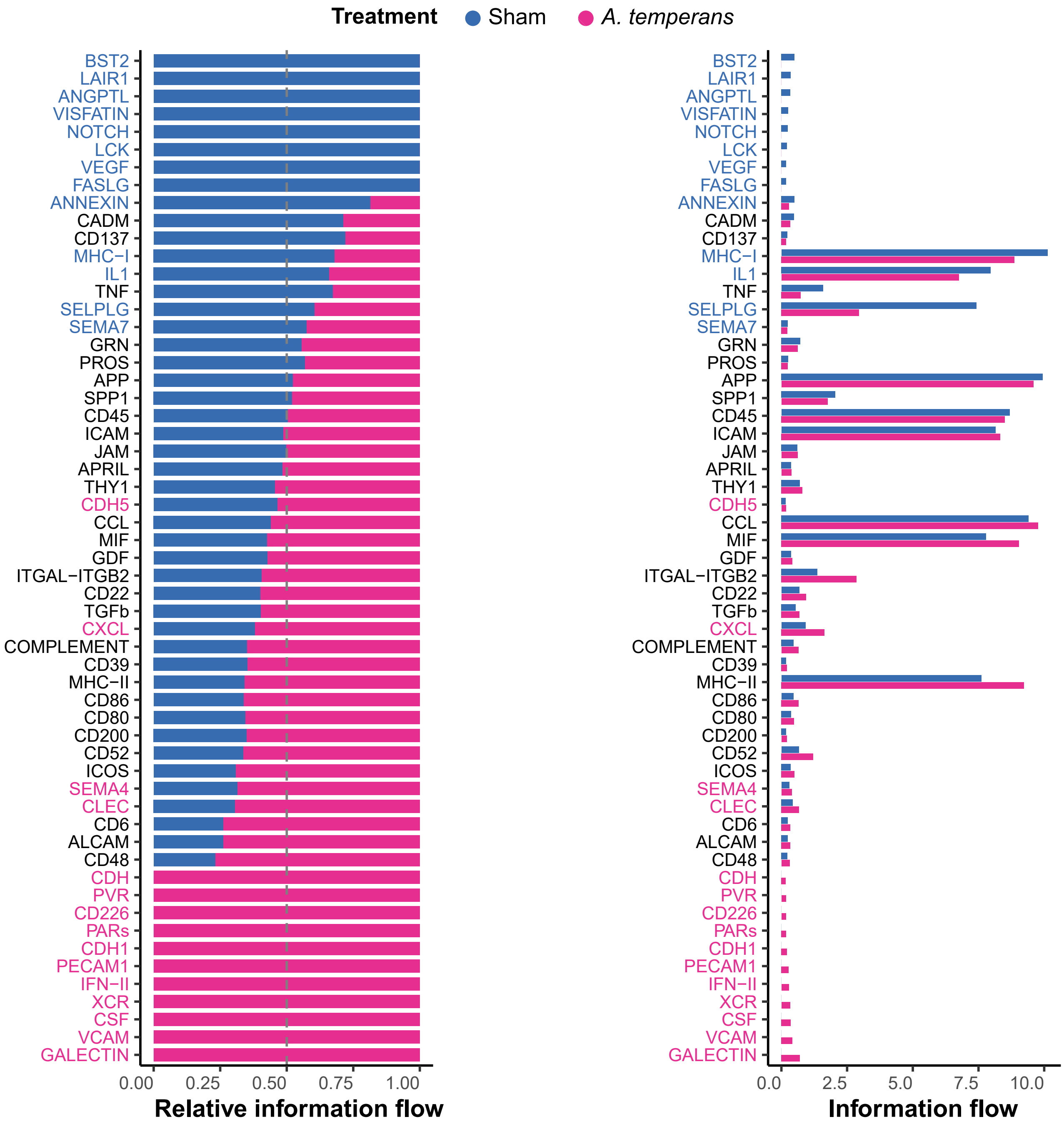

**Fig. S6 – Specificity of ligand-receptor signaling pathways by treatment group.**
Relative (left) and absolute (right) contribution of aggregate signaling pathways, ordered from sham-exclusive (top, blue) to *A. temperans*-exclusive (bottom, pink).
